# Chromatin landscape of budding yeast acquiring H3K9 methylation and its reader molecule HP1

**DOI:** 10.1101/2025.09.11.675748

**Authors:** Taiki Shimizu, Kei Fukuda, Chikako Shimura, Jun-ichi Nakayama, Masaya Oki, Yoichi Shinkai

**Author notes:** Correspondence (Y.S.).

## Abstract

Histone H3 lysine 9 (H3K9) methylation and heterochromatin protein 1 (HP1) are well conserved heterochromatin epigenome and its reader molecule. However, the details of how importance of them in heterochromatin formation still remain unclear. One of the reasons is the redundancy aspects, as there are multiple H3K9 methylation reader molecules, including HP1, and HP1 itself functions as a hub recruiting various effector molecules. To overcome those issues, we took a synthetic biology approach and introduced H3K9 methylation and HP1 into budding yeast *Saccharomyces cerevisiae*, which does not have this system, and examined its impact on transcription and chromatin compaction. We observed that the mammalian H3K9 methyltransferase can induce genome-wide H3K9 di- and tri-methylation (H3K9me2,3) in the budding yeast which mainly in the gene body region, and HP1 accumulates over the H3K9 methylated regions. The forced expression of H3K9 methyltransferase and HP1 had little impact on transcription. Furthermore, Micro-C-seq analysis revealed no significant effects on the chromatin 3D structure. These results suggest that although H3K9 methylation and recruitment of HP1 play essential roles in the epigenetic regulation of heterochromatin, they alone are not sufficient to alter the higher-order chromatin structure, at least in the gene body regions in budding yeast.

## Introduction

In eukaryotes, the genome is packed within the nucleus in the form of chromatin. The DNA is wrapped around the histone octamer which comprises of two each of H2A, H2B, H3, and H4. The chromatin density is not uniform in the nucleus, and transcription is regulated by changing chromatin densities in each genomic region. Genes present within the low-condensed regions or euchromatin are actively transcribed, whereas those present in the highly condensed regions or heterochromatin are generally repressed. Heterochromatin also contains repetitive DNA and is often found at telomeres and pericentromeres (Allshire, & Madhani, 2018; Janssen, Colmenares, & Karpen, 2018).

Transcription can be regulated by the chemical modifications of DNA and histones, known as the epigenome. DNA methylation leads to transcriptional silencing whereas histone acetylation enhances transcriptional activity (Shvedunova, & Akhtar, 2022; Smith, & Meissner, 2013). Histone methylation regulates the recruitment of reader and effector molecules, thereby, influencing transcription (Greer, & Shi, 2012). For example, the trimethylation of histone H3 lysine 4 (H3K4me3) is well correlated with transcriptional activation, whereas H3K9me2/3 and H3K27me3 are correlated with transcriptional silencing (Jambhekar, Dhall, & Shi, 2019).

H3K9 methylation and its silencing effect are widely conserved from fission yeast to humans although this system is not present in budding yeast. In mammals, multiple lysine methyltransferases regulate H3K9 methylation (Montavon, Shukeir, Erikson et al., 2021; Padeken, Methot, & Gasser, 2022). H3K9me3 is mostly regulated by SETDB1 and two SUV39H family proteins, SUV39H1 and SUV39H2 (SUV39Hs). H3K9me2 in the euchromatic regions is primarily regulated by G9a/GLP and SETDB1, whereas H3K9me2 in heterochromatic regions is controlled by these five enzymes (Fukuda, Shimura, Miura et al., 2021). SETDB1 plays a crucial role in the transcriptional silencing of transposons and heterochromatin formation at the telomere (Fukuda, & Shinkai, 2020; Gauchier, Kan, Barral et al., 2019; Kato, Takemoto, & Shinkai, 2018). SUV39Hs are critical for heterochromatin formation of telomeres, pericentromeres and retrotransposons, such as LINE-1 (Bulut-Karslioglu, De La Rosa-Velazquez, Ramirez et al., 2014; Peters, O’Carroll, Scherthan et al., 2001; Schoeftner, & Blasco, 2009). H3K9me2, catalyzed by the G9a/GLP complex, also contributes to gene and retrotransposon silencing (Padeken et al., 2022; Shinkai, & Tachibana, 2011). Although G9a and GLP alone exhibit sufficient H3K9 methyltransferase activity *in vitro*, their intracellular H3K9 methylation is maintained primarily carried out by the G9a/GLP heteromeric complex (Tachibana, Ueda, Fukuda et al., 2005).

Heterochromatin protein 1 (HP1) is a well conserved reader/effector molecule of H3K9 methylation. In mammalian cells, there are three HP1 paralogs (HP1α, β, γ). HP1 colocalizes with DAPI-dense heterochromatin regions (Zenk, Zhan, Kos et al., 2021) and contains two evolutionally conserved domains, chromodomain (CD) and chromoshadow domain (CSD). CD binds to methylated H3K9 whereas CSD is required for the dimerization of HP1 and for the interaction with HP1-binding proteins through their PxVxL motifs (Kumar, & Kono, 2020). HP1 directly induces chromatin condensation, and depletion of HP1 leads to conformational changes in mammalian chromatin (Hiragami-Hamada, Soeroes, Nikolov et al., 2016; Kilic, Felekyan, Doroshenko et al., 2018). Direct tethering of HP1 to specific gene locus in flies and mammalian cells leads to their transcriptional silencing (Gao, Han, Shang et al., 2021; Hathaway, Bell, Hodges et al., 2012; Seum, Spierer, Delattre et al., 2000). Thus, it has been suggested that H3K9 methylation-dependent gene silencing occurs, at least partly, due to chromatin condensation, which is facilitated by the interaction between H3K9me2/3-containing nucleosomes and HP1 proteins. However, the detailed role of HP1 remains unclear and it remained poorly understood whether chromatin condensation is essential for transcriptional repression. There are several proteins that can interact with HP1 and methylated H3K9 *in vivo* (Bua, Kuo, Cheung et al., 2009; Nozawa, Nagao, Masuda et al., 2010). Although the knockout/knockdown of specific proteins disrupts the interaction of every downstream factor, other proteins can compensate for deleted proteins. Therefore, it is difficult to determine the extent to which H3K9 methylation and HP1 recruitment are important for gene silencing and heterochromatin formation. To overcome this problem, we adopted a synthetic biological approach.

In the budding yeast *Saccharomyces cerevisiae*, the H3K9 methylation/HP1 reading-silencing system is absent and there are no alternate silencing systems that function via DNA and H3K27 methylation (Buhler, & Gasser, 2009; Oh, Yeom, Park et al., 2022; Rusche, Kirchmaier, & Rine, 2003). The heterochromatin regions are restricted to telomeres, rDNA, and mating type loci (HML and HMR) in *S. cerevisiae*, and the formation of these heterochromatin regions is dependent on the Sir complex (Sir2, Sir3, and Sir4) (Kueng, Oppikofer, & Gasser, 2013). Considering the simple transcriptional regulatory mechanism and alternate heterochromatin formation mechanisms, we tried to perform *de novo* H3K9 methylation-dependent gene silencing in *S. cerevisiae* to elucidate the functions of H3K9 methylation and HP1. We also investigated the role of H3K9 methylation and HP1 in regulating transcription and chromatin conformation in *S. cerevisiae*.

## Results

### Introduction of H3K9 methylation and its reader molecule HP1 in *S. cerevisiae*

First, we examined whether forced expression of mammalian H3K9 methyltransferases in budding yeast induces H3K9 methylation. To address this, we overexpressed mouse SETDB1, SUV39H1, G9A, and GLP in *S. cerevisiae*. Results showed that forced expression of SETDB1 had no effect on H3K9 methylation (H3K9me2) (Figure 1A). SETDB1 requires the associated factor ATF7IP for nuclear localization and it is known that SETDB1 ubiquitination is important for its enzymatic activity (Ishimoto, Kawamata, Uchihara et al., 2016; Sun, & Fang, 2016; Tsusaka, Shimura, & Shinkai, 2019). We co-expressed ATF7IP or UBE2E1, an E2 ubiquitin ligase involved in the ubiquitination of SETDB1, or both, with SETDB1 (Figure 1A and B). Results showed that when expressed alone, SETDB1 was mainly localized in the cytoplasm, whereas when co-expressed with ATF7IP, SETDB1 was enriched in the DAPI-positive nuclei although no H3K9 methylation was observed. When co-expressed with UBE2E1, SETDB1 showed slower mobility in SDS-PAGE (Figure 1A), indicating that the protein is ubiquitinated (Tsusaka et al., 2019) ; however, no nuclear enrichment and H3K9me2 marks were observed. However, in yeast co-expressing SETDB1, ATF7IP, and UBE2E1, H3K9me2 marks were detected. These results indicate that the subcellular localization and enzymatic activity of SETDB1 can be regulated by the co-expressing known regulatory factors. Furthermore, SETDB1 can induce H3K9 methylation in budding yeast, which does not have the H3K9 methylation system. Moreover, forced expression of G9A, GLP, or SUV39H1 alone was sufficient to induce H3K9me2 in *S. cerevisiae* (Figure 1C). Besides, these enzymes were found to be enriched in the nucleus of the budding yeast (Figure 1D).

**Fig.1.**
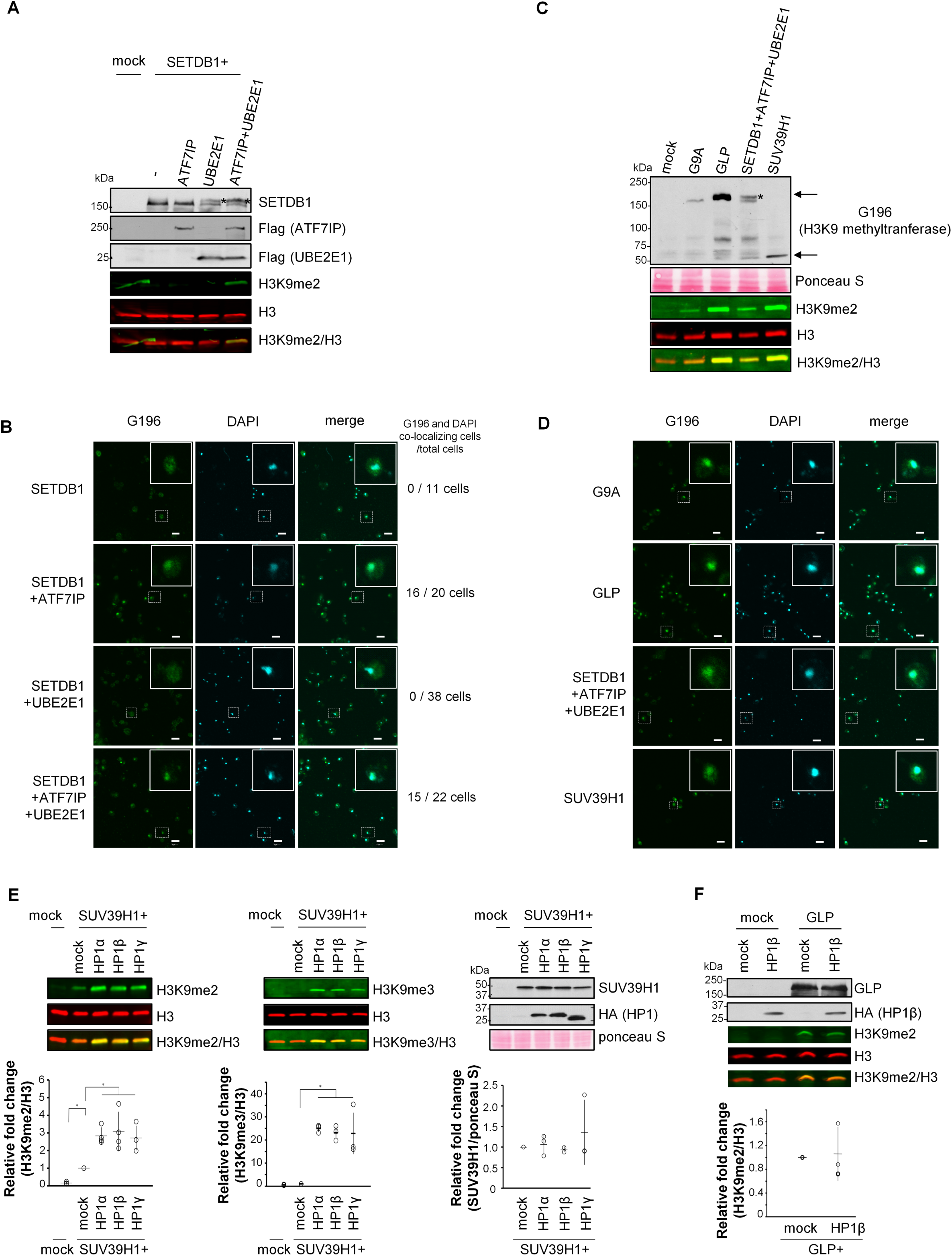
The activity of H3K9 methyltransferases in *S. Cerevisiae.* (A) G196-tagged SETDB1 (G196-SETDB1) expressing w303a strain co-expressed with Flag-ATF7IP, Flag-UBE2E1 or both Flag-ATF7IP and Flag-UBE2E1 were lysed, or their nuclei were extracted. The protein expressions and H3K9 methylation were confirmed by western blotting. The molecular weight of SETDB1 was shifted by UBE2E1 mediated mono-ubiquitination (*). Mock indicates *G196 empty vector* integrated strain. (B) The localization of SETDB1 were confirmed by immunostaining using anti-G196 antibody. Scale bar indicates 5 mm. An enlarged image of the cell circled by square is shown in the upper right corner of each panel. (C) G196-G9a, G196-GLP, G196-SETDB1 (with Flag-ATF7IP and Flag-UBE2E1), and G196-SUV39H1 expressing w303a strains grown on YPD plate were lysed in sample buffer or their nuclei were extracted. The expression of each H3K9 methyltransferase and H3K9 methylation were confirmed by western blotting. The upper and lower arrow indicate the position of G9A, GLP or SETDB1 and SUV39H1, respectively. (D) The localization of G196-G9a, G196-GLP, G196-SETDB1 co-expressed with Flag-ATF7IP and Flag-UBE2E1, and G196-SUV39H1 were confirmed by immunostaining using anti-G196 antibody. Scale bar indicates 5 mm. (E) HA-HP1a, HA-HP1b, or HA-HP1g was co-expressed in G196-SUV39H1 expressing w303a strain. After 2 days culture on YPD plate, cells were lysed, or their nuclei were extracted. Western blotting was performed using anti-H3K9me2 (left), anti-H3K9me3 (middle), and anti-SUV39H1 (right) antibodies. The signal intensity was normalized by H3 or ponceau S staining. Error bars indicate S.D. from three or four independent experiments. *p<0.05 by Tukey Kramer’s test. (F) G196mock/HAmock, G196mock/HA-HP1b, G196-GLP/HA-mock, or G196-GLP/HA-HP1b expressing w303a strain were cultured on YPD plate for 2 days. Cells were lysed, and western blotting was performed. The signal intensity of H3K9me2 was normalized by H3. Error bars indicate S.D. from three independent experiments. Statistical analysis was performed by Student t-test.

Next, we co-expressed HP1 and examined its effects on H3K9 methyltransferase activity in the budding yeast (Figure 1E). SUV39H1 forms a complex with HP1 in mammalian cells, and HP1 plays a crucial role in SUV39H-mediated H3K9me3 formation in the pericentromeric region (Melcher, Schmid, Aagaard et al., 2000; Muramatsu, Kimura, Kotoshiba et al., 2016; Yamamoto, & Sonoda, 2003). Co-expression of SUV39H1 with either HP1α, β or γ enhanced H3K9 methyltransferase activity, enhanced the H3K9me2 signal and almost undetectable H3K9me3 was also detected, even though the amount of SUV39H1 was unchanged (Figure 1E). However, when GLP was co-expressed with HP1β, the enzymatic activity of GLP remained unchanged (Figure 1F).

### Global H3K9 methylation and HP1β distribution on yeast chromatin

Since H3K9 methylation was induced in yeast strains expressing SUV39H1 (Figure 1C) and this methylation was further enhanced by co-expression of HP1 (Figure 1E), we further characterized the genome-wide distribution of H3K9 methylation in strains expressing SUV39H1 either alone or with HP1β. We analyzed the genome-wide distribution of H3K9me2 marks in cells expressing SUV39H1 alone, and H3K9me2 marks, H3K9me3 marks, and HP1β localization in cells expressing SUV39H1 and HP1β (Figure 2A, B). As shown in Figure 2A, SUV39H1 expression alone induced the formation of H3K9me2 marks, mainly in the gene body regions. When co-expressed with HP1β, the H3K9me2 marks remained in the gene body region, but their signal intensity was enhanced, which is in agreement with the results of western blot (Figure S1A). The enrichments of H3K9me3 and HP1 β in cells expressing SUV39H1 and HP1β were highly correlated with that of H3K9me2 (Spearman correlation values: 0.81–0.89 with H3K9me2 and 0.90–0.94 with H3K9me3) (Figure 2C and Figure S1B). Similar to the distribution of H3K9me2 and H3K9me3 marks, HP1β enrichment was very low near regions around -0.1 kb from the TSS (Figure 2A). We validated ChIP-seq data analysis for H3K9me2, me3 and HP1β distribution and their accumulation in some enriched genes by ChIP-qPCR. Compared to regions upstream of TSS, H3K9me2 and H3K9me3 signals were more prominent in the gene body regions, and these signals were further enhanced by HP1β co-expression (Figure 2D). We also examined the effects of SUV39H1 and HP1 co-expression on other histone modifications, such as H3K9ac, H3K4me3, and H3K27ac by western blotting (Figure S2). It was not significant difference, but H3K9 acetylation tended to be decreased following HP1 co-expression. Other histone modifications remained unaffected.

**Fig. 2.**
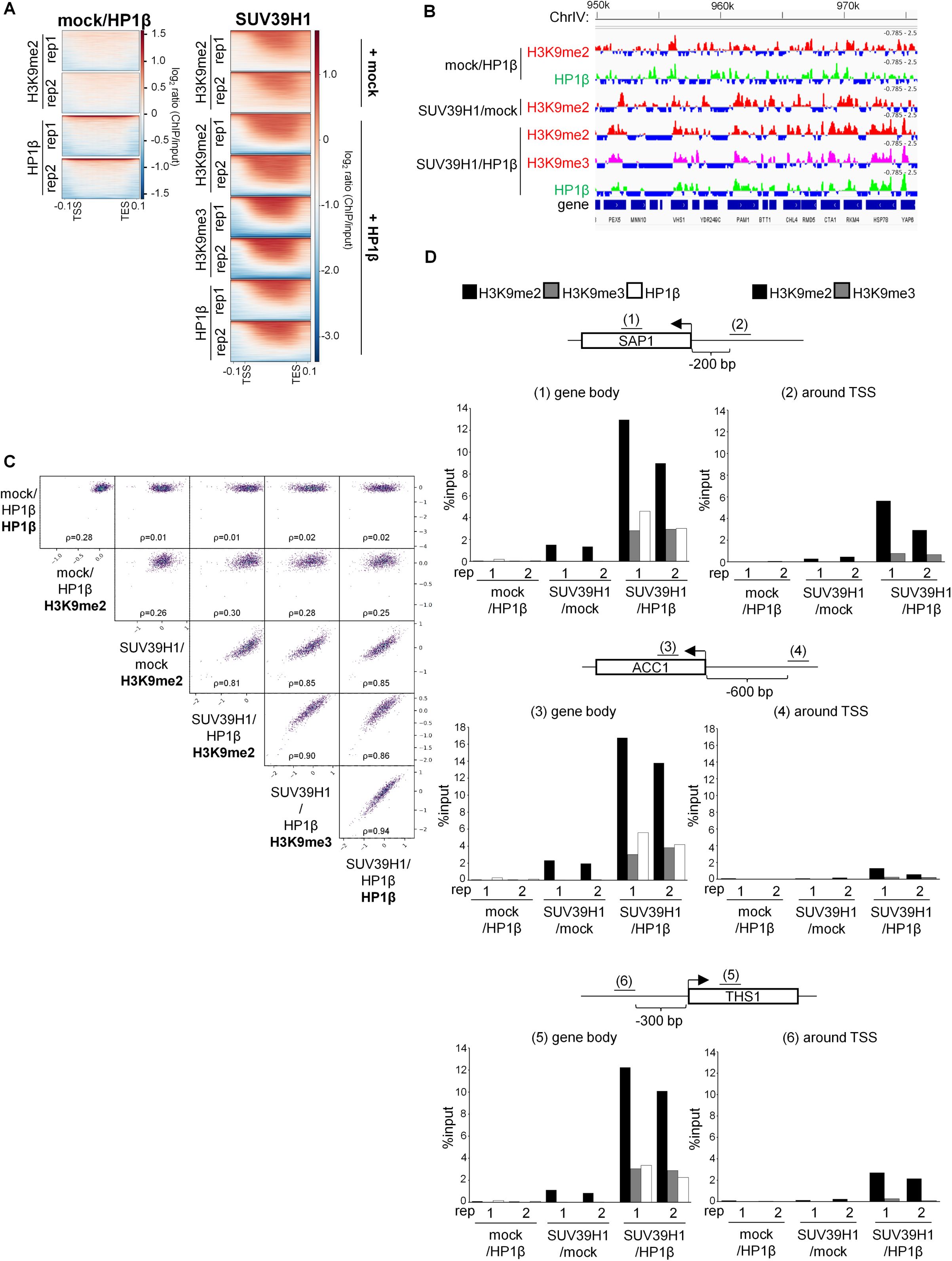
The effect of HP1 on H3K9 methylation by SUV39H1. (A) Heatmaps for localization of H3K9me2, H3K9me3, and HP1b in G196-SUV39H1 strains co-expressed with or without HA-HP1b. H3K9me2 ChIP samples were obtained from mock/HP1b strain, SUV39H1/mock strain and SUV39H1/HP1b strain. H3K9me3 ChIP sample was obtained from SUV39H1/HP1b strain. HP1b ChIP samples were obtained from mock/HP1b strain and SUV39H1/HP1b strain. (B) ChIP-seq data of replication 1 in chromosome IV 950kb-975kb was viewed in Integrative Genomics Viewer. H3K9me2 or H3K9me3 were in red. HP1b were in green. sacCer3 was used for reference gene. (C) Spearman correlation map of H3K9me2, H3K9me3, and HP1b ChIP-seq data of replication 1 samples. (D) ChIP qPCR was performed to determine the quantity of H3K9 methylation and HP1 in some gene bodies (*SAP1*, *THS1*, *ACC1*) and the quantity of H3K9 methylation at the intergenic regions (pre*SAP1*, pre*THS1*, pre*ACC1*). Both replication 1 and 2 were shown.

As described in the introduction section, the H3K9 methyltransferase-HP1 write-read system is absent in budding yeast, although H3K36 methylation is present. One of the H3K36 demethylases, Rph1, can demethylate H3K9 if expressed in mammalian cells (Klose et al., 2007). To test the possibility that Rph1 eliminates SUV39H1-mediated H3K9 methylation, especially around the TSS in yeast, we examined the levels and distribution of H3K9 methylation following *RPH1* inactivation in strains expressing SUV39H1 and HP1β (Figure S3A). Western blot and ChIP-qPCR analyses showed no clear enhancement of H3K9me2 and H3K9me2 levels around the TSS of the analyzed genes (Figure S3B,C), indicating that poor H3K9 methylation around TSSs in strains expressing SUV39H1 with/without HP1β is independent of Rph1.

### The effect of HP1β on global transcription and chromatin condensation

Next, we examined the impact of H3K9 methyltransferase and HP1 expression on yeast transcription using RNA-seq analysis. We found that forced expression of SUV39H1 affected the levels of only six genes (five upregulated and one downregulated; FDR < 0.05) (Figure 3A, blue marked dots). Furthermore, co-expression of HP1β in SUV39H1 expressing strain affected the transcription of only 15 genes (5 upregulated and 10 downregulated) (Figure 3A and Figure S4A). Distribution and intensity of H3K9me2/3 around the genes affected by SUV39H1 or SUV39H1+HP1β expression were shown (Figure S4B), but no clear feature was observed between the affected genes and unaffected their neighboring genes. These results indicate that forced expression of SUV39H1 induces sequence-independent genome-wide formation of H3K9me2 or H3K9me3 marks within the gene body regions and co-expression of HP1β increases the methylation levels. Moreover, HP1β localizes to these methylated regions, although it has no impact on yeast gene transcription.

**Fig. 3.**
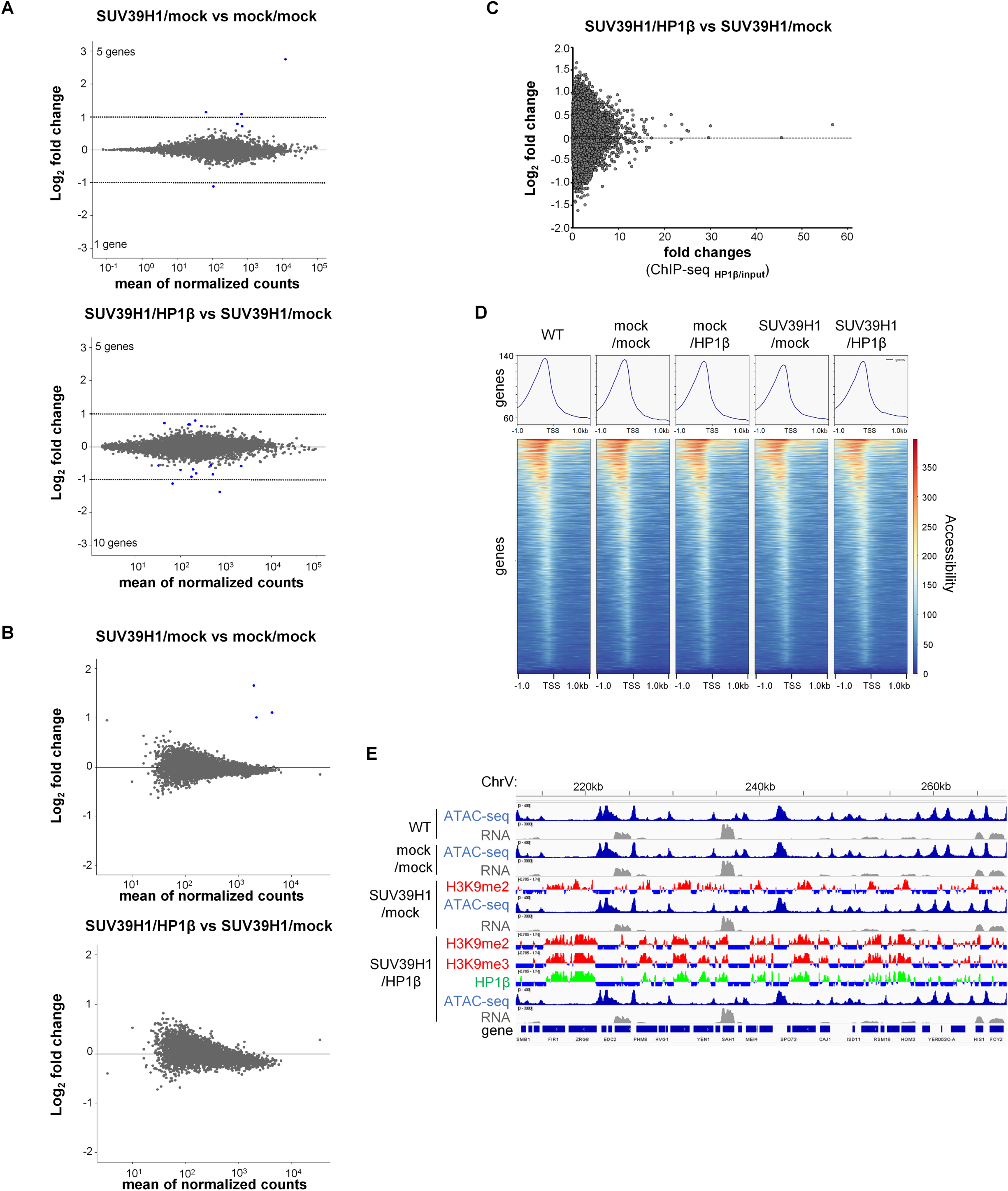
The effect of SUV39H1 and HP1b expression on gene expression and chromatin compaction in budding yeast. (A) Gene expression changes by forced expression of G196-SUV39H1 and by co-expression of HA-HP1b in SUV39H1 expressing strain were analyzed. The differentially expressed genes (FDR<0.05) between mock (mock/mock) vs SUV39H1 expressing (SUV39H1/mock) strain (upper) and SUV39H1/mock vs SUV39H1+ HP1b expressing (SUV39H1/ HP1b) strain (lower panel) were indicated as blue. (B) The number of sequence reads in each ATAC-seq peak was counted, and the difference of the number between SUV39H1/mock vs mock/mock (upper) or SUV39h1/ HP1b vs SUV39H1/mock (lower) was compared. The different peak counts regions (FDR<0.05) were indicated in blue dots. (C) Normalized read numbers of ATAC-seq peaks in every 100 bp of whole genome of the strain expressing SUV39H1 (SUV39H1/mock) and SUV39H1+ HP1b (SUV39H1/HP1b) were compared. The ratio was plotted with the HP1b ChIP-seq enrichment levels in the same 100 bp regions of SUV39H1/HP1b strain. X-axis indicates the amount of HP1b, and Y-axis indicates the ratio of normalized read number of ATAC-seq peaks of two strains. The analysis data using replication 1 was shown. (D) Heatmaps for the localization of each ATAC-seq signal using mapping data. Replication 1 data were shown. (E) RNA-seq and ATAC-seq tracks with ChIP-seq of replication 1 in chromosome V were viewed in Integrative Genomics Viewer. ATAC-seq peaks were in blue. RNA-seq was in gray. H3K9me2 was in red. HP1b was in green.

However, it was not clear from previous analyses how much amount of H3K9 methylation was induced globally. Therefore, we compared the H3K9me2 and H3K9me3 levels in these yeast stains with those in mouse ES cells (mESCs) (Figure S5). Results showed that H3K9me2 levels in yeast expressing only SUV39H1 were approximately the same as that in mESCs, whereas in lines expressing SUV39H1 and HP1β, the levels of H3K9me2 were 3 to 10 times higher than that in mESCs and H3K9me3 levels were also found to be several times higher. These results suggest that the global H3K9me2 and H3K9me3 levels in yeast expressing SUV39H1 and HP1β are comparable to those of mESCs.

As HP1 condenses chromatin by bridging two methylated nucleosomes or LLPS, next we performed ATAC-seq to investigate whether co-expression of SUV39H1 and HP1β alters chromatin condensation sates in *S.* cerevisiae. As shown in Figure 3B and Figure S6, all ATAC-seq data showed highly correlated. Furthermore, there was no correlation between the differential signal intensities of ATAC-seq between SUV39H1 alone and SUV39H1+HP1β dual-expressing strains and the HP1β enrichment in the dual-expressing strains (Figure 3C). The signals of ATAC-seq of all strains examined were enriched in front of TSS (Figure 3D), and a comparison of ATAC-seq and ChIP-seq data showed that ATAC-seq signals and H3K9me2, H3K9me3, and HP1β sites were segregated over the genome, as shown in Figure 3E. We concluded that these epigenetic modifications did not affect chromatin accessibility.

Next, we asked why H3K9me2 was not induced around the TSS when SUV39H1 was expressed. A possible reason is that most genes in yeast are not heterochromatinized and transcribed, therefore, lysine residues of H3 and H4 around the TSS are highly acetylated, including H3K9. Thus, methylation cannot be further induced at H3K9 residues in these regions. Based on RNA-seq analysis of the wild-type (WT) strain analyzed in this study, we divided genes into three groups: very low expression group (<5 transcripts per million (TPM)), intermediate expression group (5– 80 TPM), and high expression group (>80 TPM), and re-analyzed the distribution of H3K9me2 in SUV39H1 and HP1β expressing strains. Results showed that H3K9me2 levels in the gene body and around the TSS were similar in the very low expression group. The higher the expression level, the lower H3K9me2 was observed around the TSS (Figure S7A). Next, we examined the acetylation pattern of H3 lysine residues (K9, K14, K18, and K27) and H3K36me3 in genes of these three groups using the publicly available genome-wide ChIP-seq data of budding yeasts (Weiner, Hsieh, Appleboim et al., 2015), Figure S7B,. Results showed that the acetylation of lysine residues, especially K9, K14, and K18 in H3 was negatively correlated with H3K9me2. Furthermore, the distribution of H3K36me3 marks was well correlated with that of H3K9me2 in intermediate and highly expressed genes, but not in genes with a low expression (since H3K36me3 is a transcription-linked and induced modification). It is possible that either H3K9 hyperacetylation prevents methylation at these sites or hyperacetylation of all H3 lysine residues, including H3K9 and K14, comprehensively suppresses SUV39H1-mediated H3K9 methylation, as reported in fission yeast (Nakayama, Rice, Strahl et al., 2001; Oya, Nakagawa, Yoshimura et al., 2019). Furthermore, chromatin accessibility around TSS of SUV39H1 either alone or in the presence of HP1β showed no significant difference in the three groups (Figure S8).

### The effect of HP1β expression on genome conformation

ATAC-seq analysis showed that the expression of H3K9 methyltransferase and its reader molecule, HP1, had no obvious effect on genome-wide chromatin accessibility. However, in organisms possessing H3K9 methylation regulatory systems, H3K9 methylation-mediated HP1 recruitment/binding to chromatin contributes to heterochromatin formation (Hiragami-Hamada et al., 2016; Lehnertz, Ueda, Derijck et al., 2003; Nakayama et al., 2001; Rusche et al., 2003). Therefore, we analyzed the chromatin higher-order structure in budding yeast expressing H3K9 methyltransferase SUV39H1 and HP1β using Micro-C XL analysis (Hsieh, Fudenberg, Goloborodko et al., 2016). We analyzed the relationship between contact frequency and genome distance at 10 kb resolution. Results showed that the WT, SUV39H1/mock, and SUV39H1/HP1β strains had similar genome contact frequencies (Figure 4A, Figure S9A). We also compared the correlation of TAD insulation scores of each 100 bp matrix, and results showed a high correlation among different strains (WT-SUV39H1/mock: Spearman = 0.95, WT-SUV39H1/HP1β: Spearman = 0.94, SUV39H1/mock-SUV39H1/HP1β: Spearman = 0.97) (Figure 4B, Figure S9B). Figure 4C shows contact heatmaps with TAD insulation scores and localization of HP1β in a local region (chromosome III 210–310 kb) at a resolution of 100 bp. The contact pattern and TAD regions remained unchanged following SUV39H1 and HP1β expression (Figure 4C, Figure S9C). We also compared the contact heatmap in the global region (chromosomes I, II, III, and IV) at 10 kb resolution, and again no clear differences were observed (Figure 4D). The telomere-telomere and centromere-centromere interactions were not affected by SUV39H1 and HP1 expression. These results suggest that H3K9 methylation and HP1β recruitment are not sufficient to alter the 3D genome organization in budding yeast.

**Fig.4.**
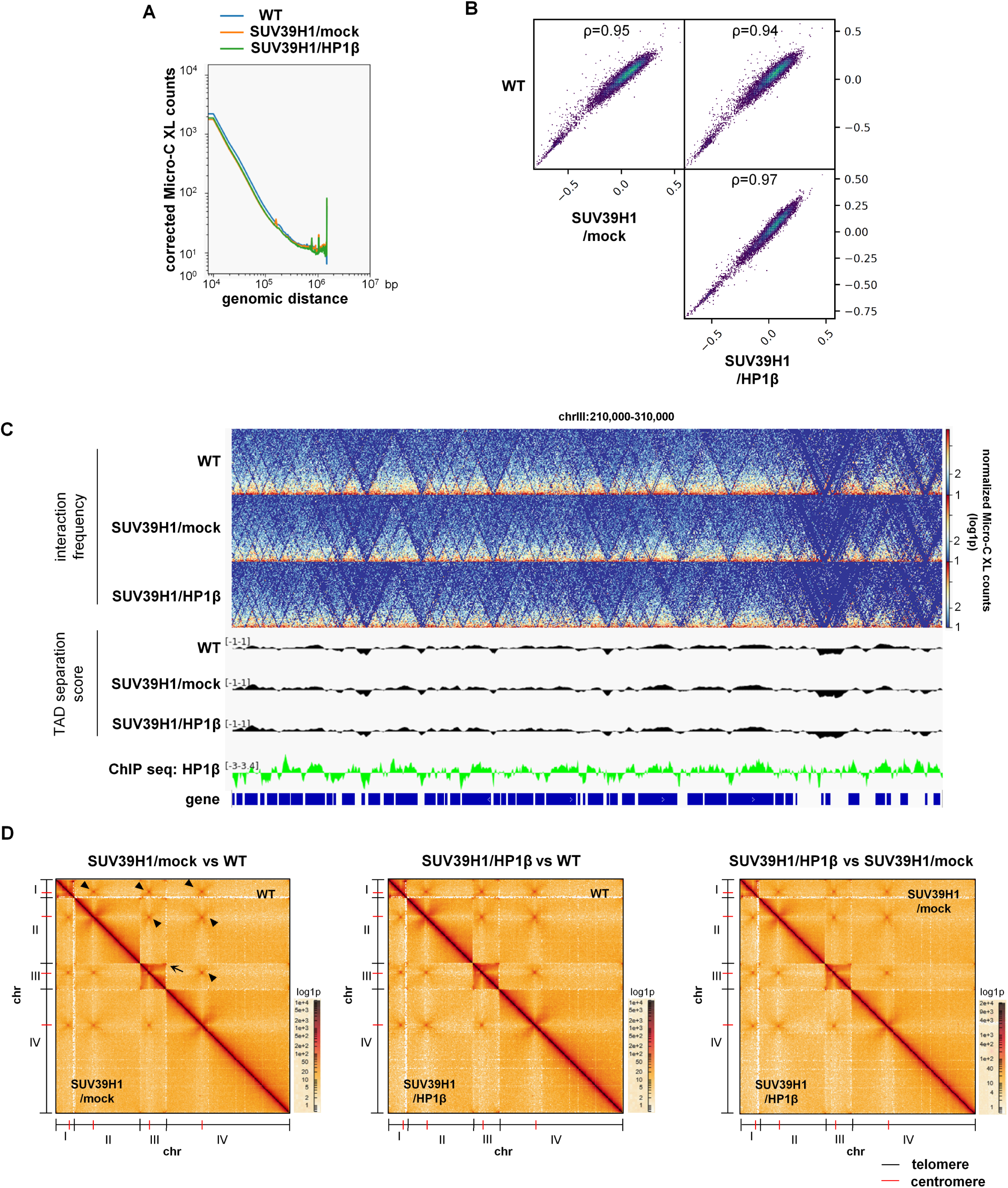
The effect of HP1β on genome conformation in SUV39H1 expressing strain. (A) Genome contact frequency in WT, SUV39H1/mock, and SUV39H1/HP1b strains were compared using 10k bp windows. (B) TAD regions in 100 bp bin contact matrix was identified, and the spearman correlation was plotted. (C) Hi-C matrix and TAD domain score were plotted with HP1b chip-seq data derived from SUV39H1/HP1b replication 1. Chromosome III (210,000-310,000 bp) was shown. Separation score indicates the mean z-score of all the matrix contacts between the left and right regions. (D) Genome wide contact heatmap in chromosome I, II, III, and IV at 10kbp resolution. Arrows indicate centromere interactions, and arrow heads indicate telomere interactions.

### The tethering of H3K9 methyltransferases and recruitment of HP1 to *a1* gene and its impact on transcription

Our results showed that forced expression of H3K9 methyltransferase and HP1β had no effect on transcription in budding yeast. However, since H3K9 methylation was relatively low around TSS, and H3K9 methylation in the promoter region around TSS is important for transcriptional regulation, we performed experiments to enhance H3K9 methylation specifically around the TSS.

We tethered H3K9 methyltransferases directly to a specific locus using the GAL4/UAS system. The mating type “α” strain ROY2042 contains 4×UAS sequences upstream of the reporter transgene *a1* near the *URA3* gene locus which determines the mating type of *S. cerevisiae*. Another strain ROY2041 was used as a control which also possesses the *a1* transgene but without the UAS sequences (Oki, Valenzuela, Chiba et al., 2004) (Figure 5A). We deleted the *SIR3* gene to disrupt heterochromatin formation because *a1* is located in the heterochromatin region and is permanently silenced in the *SIR3* intact strains. Each GAL4-binding domain (GBD)-H3K9 methyltransferase fusion molecule was expressed in these strains and H3K9 methylation at the *a1* gene was determined by ChIP-qPCR. Results showed that although the methylation level was low, H3K9me2 was significantly induced in the body of the *a1* gene (-0.7 kb from 4×UAS) in every ROY2042 *Δsir3* strain expressing different GBD-H3K9 methyltransferases (plus ATF7IP and UBF2E1 for SETDB1) but not in ROY2041 *Δsir3* strains (Figure 5B-E). We also checked the H3K9 methylation status around the UAS by qPCR using the GBD-SUUV39H1 strain. H3K9me2 marks were more prominent at the -0.2 kb position from the 4×UAS site compared to those at the *a1* gene body region (-0.7 kb) in ROY2042 *Δsir3* strain, but no accumulation of H3K9me2 marks was detected at further up or downstream regions (-3, +2.4 and +3 kb) from the 4×UAS site (Figure 5F).

**Fig.5.**
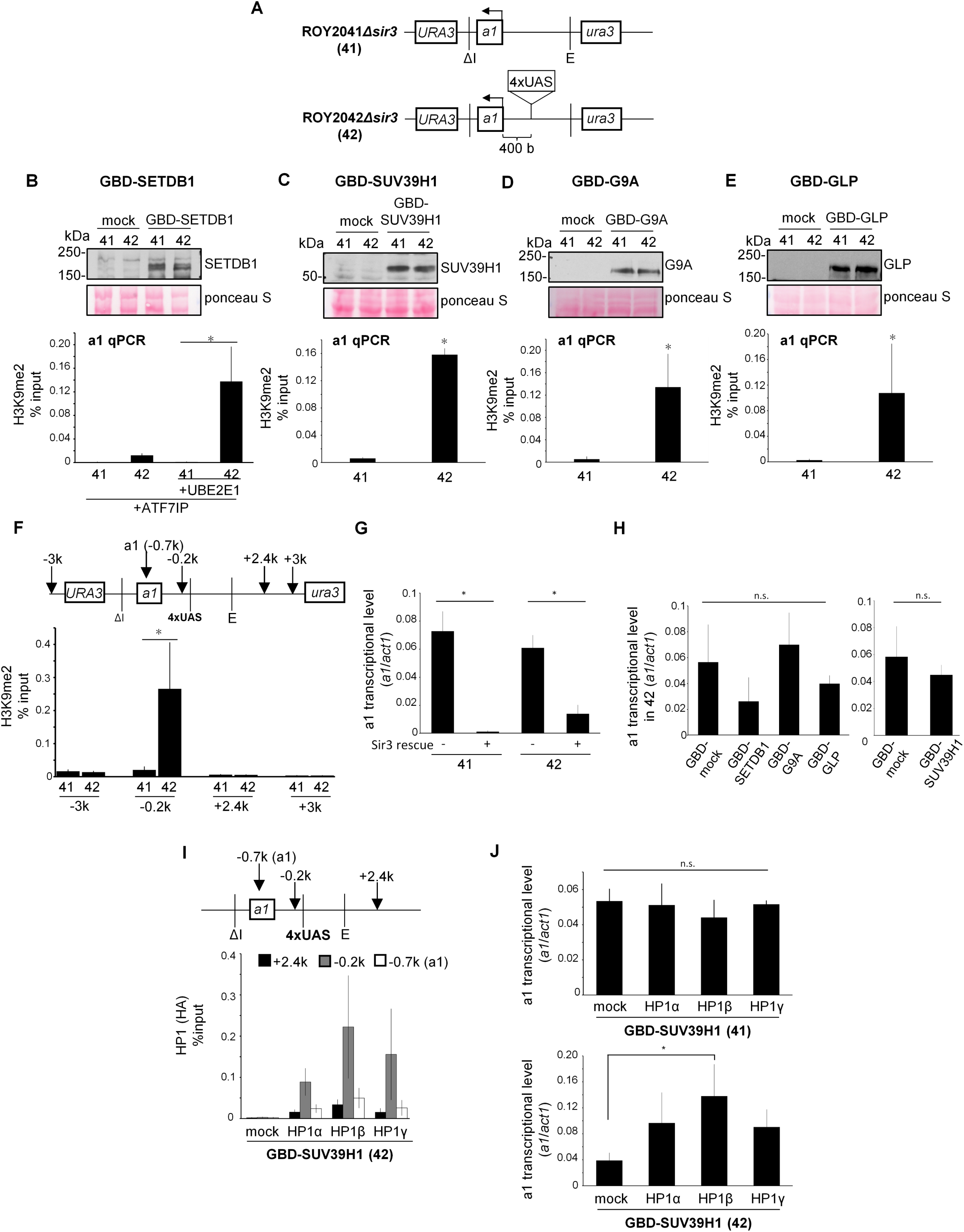
Tethering GBD-H3K9 methyltransferases at *a1* gene. (A) *HMR* sequence containing plasmid was integrated into *URA3* locus of *hmr* deletion mutant (ROY2041). ROY2042 strain the same as ROY2041 strain except for containing 4xUAS sequences in the upstream of *a1* gene. The length between UAS and the start codon of *a1* gene was about 400 bp. *SIR3* gene was deleted in both strains. (B-E) Gal4 binding domain (GBD) fusion H3K9 methyltransferases were integrated into the genome of ROY2041*Δsir3* (-UAS) or ROY2042*Δsir3* strain (+UAS). Cells were cultured in YPD medium, and chromatin immunoprecipitation was performed using anti-H3K9me2. H3K9 methylation in *a1* gene was confirmed by qPCR with the primers targeting *a1* gene body (about 700 bp away from UAS). Error bars indicate S.D. from three independent experiments. *p<0.05 by Tukey Kramer’s test or student’s t test. (F) H3K9 di-methylation status around 4xUAS sequences of GBD-SUV39H1 expressing strains was confirmed using the same ChIP sample of Fig 2C. The numbers (-3k, -0.2k, +2.4k, +3k) indicate the approximate distance from 4xUAS sequences. *p<0.05 by student’s t test. (G) mRNA was purified from ROY2041Δsir3 (-(-UAS)), ROY2041*Δsir3*+Sir3 rescue (Sir3(-UAS)), ROY2042Δsir3 (+(UAS)), and ROY2042*Δsir3*+Sir3 (Sir3(+UAS) rescue strains. *a1* transcriptional level was quantified by qPCR. *ACT1* was quantified for normalization. Error bars indicate S.D. from four independent experiments. *p<0.05 by Tukey Kramer’s test. (H) mRNA was purified from each H3K9 methyltransferase expressing strain, and *a1* transcriptional level was quantified by qPCR. *ACT1* was quantified for normalization. Mock indicates GBD empty vector, and GBD-SETDB1 strain was co-expressed with ATF7IP and UBE2E1. Error bars indicate S.D. from three independent experiments. Statistical analysis was performed by Tukey Kramer’s test. n.s. means no significant differences. (I) HA-empty vector, HA-HP1a, HA-HP1b, or HA-HP1g was co-expressed in GBD-SUV39H1 expressing ROY2042*Δsir3* strain. Each strain was cultured in YPD medium, and chromatin immunoprecipitation was performed using anti-HA antibody. Localization of HP1 around 4xUAS sequences was confirmed by qPCR. The numbers (-0.7k, -0.2k, +2.4k) indicate the approximate distance from 4xUAS sequences. (J) *a1* transcriptional levels in GBD-SUV39H1/HA-HP1 expressing ROY2041*Δsir3* strains (top) and ROY2042*Δsir3* (bottom) were quantified by qPCR. *ACT1* was quantified for normalization. Error bars indicate S.D. from three independent experiments. *p<0.05 by Dunnett‘s test. n.s. means no significant differences.

Next, we investigated whether H3K9 methylation affects *a1* transcription using RT-PCR. As shown in Figure 5G, the *a1* gene was de-repressed in *Δsir3* strains with or without 4×UAS, while Sir3 re-expression silenced *a1* transcription, although the silencing was less efficient in the ROY2042 strain compared to that in the ROY2041 strain that lacks 4×UAS insertion. Although *a1* expression was slightly reduced in the GBD-SETDB1 expressing strain, there were no significant differences in *a1* transcript levels between the mock and all other GBD-H3K9 methyltransferase-expressing strains (Figure 5H).

We also performed the mating assay to validate the effects of H3K9 methylation on *a1* transcription. Strains in which *a1* is suppressed mate with mating type “a” strains and grow in a specific synthetic defined medium. When GBD-H3K9 methyltransferase expressing strains were mixed with the “a” type strain, mating was not observed in any strain except for the Sir3 rescued strain (Figure S10A). This result validated our qRT-PCR observations that GBD-H3K9 methyltransferase-mediated H3K9 methylation around the TSS of the *a1* gene is unable to suppress *a1* transcription. In conclusion, H3K9 methylation alone is insufficient for suppressing *a1* transcription.

Finally, we analyzed the effect of HP1 expression on *a1* transcription in budding yeast. HP1α, HP1β, or HP1γ were co-expressed in each H3K9 methyltransferase-expressing strain (Figure S10B). In ROY2042 *Δsir3* strains expressing GBD-SUV39H1 and HP1s, HP1 enrichment at the *a1* locus (near the 4×UAS site) was confirmed by ChIP-qPCR (Figure 5I). However, *a1* transcription was increased several-fold in these strains (Figure 5J). It is known that the H3K9 methylation-HP1 pathway can induce transcriptional activation under some circumstances (Hediger, & Gasser, 2006; Vakoc, Mandat, Olenchock et al., 2005). It might be possible that H3K9 methylation and HP1 crosstalk with the transcriptional activation pathway in budding yeast. These results suggest that forced induction of H3K9 methylation and HP1 accumulation around the TSS of the *a1* gene do not suppress its transcription.

## Discussion

H3K9 methylation-mediated gene silencing and heterochromatin formation are well studied in various model organisms, such as *Schizosaccharomyces pombe*, *Drosophila melanogaster*, and mammals (Fukuda et al., 2021; Nakayama et al., 2001; Schotta, Ebert, Krauss et al., 2002; Xu, & Jiang, 2020). Furthermore, HP1 has been extensively characterized as a reader molecule for H3K9 methylation and plays an important role in gene silencing and chromatin condensation (Azzaz, Vitalini, Thomas et al., 2014; Larson, Elnatan, Keenen et al., 2017). Herein, we investigated the role of H3K9 methylation and its reader and effector molecule HP1 in transcriptional regulation and chromatin condensation using a synthetic biology approach. In this approach, mammalian H3K9 methyltransferases and HP1 were introduced into budding yeast that lacks the H3K9 methylation system.

The functional regulation of SETDB1 can be recapitulated in budding yeast. This regulation is conserved in *D. melanogaster* and *Caenorhabditis elegans*. In budding yeast, the ATF7IP paralog regulates the nuclear localization of SETDB1 (Delaney, Methot, Guidi et al., 2019; Koch, Honemann-Capito, Egger-Adam et al., 2009; Mutlu, Chen, Moresco et al., 2018), suggesting that budding yeast uses a conserved common machinery for the nuclear transport of SETDB1. As previously described, H3K9me2 marks are maintained primarily by the G9A/GLP heteromeric complex, and not by G9A or GLP alone, although each protein exhibits sufficient H3K9 methyltransferase activity *in vitro* (Tachibana et al., 2005). However, we observed that GLP alone can induce global H3K9 methylation when expressed in budding yeast. There could be several mechanistic explanations for these differences. Given the small size of the budding yeast genome, high expression of the enzyme might have ensured sufficient H3K9 methylation in cells. Moreover, the DNA replication and growth rate of yeast cells are significantly higher compared to those of mammalian cells, thus, allowing sufficient time for the free histones to encounter the methyltransferase. It might also be possible that specific mechanisms that allow G9a/GLP heterocomplex-specific methylation or inhibit methylation by G9A or GLP homodimers are absent in the budding yeast. Finally, it can be speculated that although the epigenetic information, including H3K9 methylation, is maintained following DNA replication in mammalian cells via specific mechanisms, these mechanisms are absent in budding yeast, which under normal circumstances, lacks H3K9 methylation. Therefore, the global methylation distribution induced by H3K9 methyltransferases, typically SUV39H1 observed in this study, was not strictly regulated and only around the TSS, where H3K9 acetylation might antagonize H3K9 methylation, the efficiency was low. The distribution of H3K9 and other lysine acetylation marks over genes was negatively correlated with that of H3K9 methylation induced by SUV39H1 in the presence of HP1 (Figure S7B), suggesting a critical role of H3 hyperacetylation in preventing H3K9 methylation. To better understand the transcriptional repression by H3K9 methylation, it is important to understand the mechanism by which H3K9 methylation is prevented around the TSS in budding yeast and whether this regulatory mechanism is general in eukaryotes. Our findings that HP1 enhances the methyltransferase activity of SUV39H and promotes the conversion of me2 to me3 are similar to the findings of previous studies on the mammalian pericentromeric region (Muller, Fierz, Bittova et al., 2016; Muramatsu et al., 2016). Although SUV39H is mainly targeted to the constitutive heterochromatic regions such as pericentromeres and telomeres and regulates the formation of H3K9me3 marks, no such localization of SUV39H to the heterochromatic regions was observed in budding yeast. This suggests that H3K9 methylation in budding yeast is due to the absence of specific regulatory mechanisms that are usually active in mammalian cells.

In the non-targeted system, SUV39H1-mediated H3K9 methylation mainly occurred in the gene body, while its accumulation was strongly suppressed around the TSS, therefore, its effect on transcription could not be fully understood. In the present study, little or no effect on global transcription was observed when SUV39H1 and HP1 were expressed together in budding yeast. Furthermore, no transcriptional repression was observed in the GAL4/UAS system in which H3K9 methyltransferase and HP1 were forced to recruit around the TSS of the *a1* reporter transgene. It might be possible that either sufficient H3K9 methylation was not induced, and thus, no repressive effects were observed, or accumulation of H3K9me-HP1 around the TSS alone is not sufficient to repress transcription. We speculate that further recruitment of effectors that function downstream of HP1 is necessary to suppress transcription. Indeed, similar study had been published recently that examining whether heterochromatin formation and transcriptional repression could be induced by introducing H3K9 methylation and HP1 in budding yeast (Yuan, & Moazed, 2024). In this study, they introduced human SUV39H2 SET domain fused with the bacterial tetracycline repressor TetR (TetR-SET) and Chp2, one of the two HP1 proteins in fission yeast. TetR-SET could induce H3K9me2, 3 at the *tetO_10x_-ADE2* reporter loci. When Chp2 was expressed in strains expressing TetR-SET, the *ADE2* reporter gene silencing could not be induced; however, suppression could be induced when Chp2 fused with fission yeast HDAC, Sir2 was expressed, indicating that H3K9 methylation and HP1 accumulation alone are insufficient for transcriptional repression in budding yeast.

Since the ATAC-seq signal is generally observed around the TSS, it is not surprising that the ATAC-seq signal was not attenuated by H3K9 methylation and HP1 expression. Hi-C (Micro-C) analysis also revealed no obvious changes in chromatin 3D structure following SUV39H1 and HP1 expression, which is contradictory to the findings of previous studies showing that chromatin H3K9 methylation and HP1 expression leads to chromatin condensation (Hiragami-Hamada et al., 2016; Kilic et al., 2018). We observed that the genome-wide abundance of H3K9me2 and H3K9me3 marks in our budding yeast model was comparable to that in mammalian cells, suggesting that H3K9me2 formation and HP1 expression are not sufficient to alter the 3D chromatin structure, especially chromatin condensation, which may require other downstream factors. In the future studies, we would like to examine how the 3D structure of chromatin is affected by the introduction of further downstream factors in this system.

Similar to our report, a recent study analyzed the effects of DNA methylation in budding yeast on transcription and chromatin structure using a synthetic biology approach (Buitrago, Labrador, Arcon et al., 2021). They found that inducing DNA methylation in yeast clearly impacts both transcription and chromatin structure. Their results showed that DNA methylation can affect chromatin structure and function, even in the absence of cellular machinery evolved to recognize and process this epigenome. In contrast, methylation of H3K9 in the N-terminal histone tail of nucleosomes and introduction/recruitment of its readout molecule, HP1, have no clear impact on transcription and genome organization, suggesting that H3K9 methylation-HP1 itself mainly functions as a hub for recruiting other factors of this pathway. However, it remains unclear why H3K9 methylation and HP1 are highly conserved and function as important epigenomes for heterochromatin regulation across species. The model system established in this study will be useful to address this issue in future studies.

## Experimental Procedures

### Yeast strains and plasmids construction

w303a strain (MATa *ADE2 lys2Δ his3-11 leu2-5,112 trp1-1 ura3-1*) was used for genome wide analysis. ROY2041 strain (MATα *ADE2 lys2Δ his3-11 leu2-5,112 trp1-1 hmrΔ::bgl-bclΔ ura3-1::HMRΔI::URA3*) and ROY2042 strain (MATα *ADE2 lys2Δ his3-11 leu2-5,112 trp1-1 hmrΔ::bgl-bclΔ ura3-1::HMRΔI+4xGal4 bs at mat a2(ECONI)::URA3*) (Oki et al., 2004) with *sir3* deletion (s*ir3Δ::KanMX*) were used for gene specific analysis.

N-terminal 3xG196 tagged mouse H3K9 methyltransferases (*Setdb1, Suv39h1, Ehmt2, Ehmt1*) were inserted into pRS406 vector. N-terminal 3xHA tagged mouse HP1 (*Cbx1,3,5*) were inserted into pRS405 vector. N-terminal 3xFlag tagged mouse *Atf7ip*, *Ube2e1* and *Atf7ip-2A-Ube2e1* were inserted into pRS403 vector. N-terminal GBD fused mouse H3K9 methyltransferases were inserted into pGBK-RC vector (Ito, Tashiro, Muta et al., 2000). These plasmids were integrated into genome using each selection markers (pRS406: *URA3*, pRS405: *LEU2*, pRS403: *HIS3*, pGBK-RC: *TRP1*). Every gene except for *Suv39h1* was expressed using *ADH1* promoter, and *Suv39h1* was expressed using *TPI1* promoter. *SIR3* was inserted into pRS403 vector with *SIR3* promoter. In-Fusion HD Cloning Kit (Takara Bio Inc.) was used for plasmids construction.

### Culture conditions

For western blot analysis, yeast cells grown on YPD plate for 2 days were used because H3K9 methylation was not detected by western blot from the yeast cultured in YPD medium. For next generation sequencing analysis, yeast cells grown on YPD plate for 2 days followed by cultured in YPD medium for 3 hours were used in order to make the digestion of cell walls easier. The cells for RNA-seq, ATAC-seq, and Micro-C XL were prepared from same culture except for RNA-seq of Suv39h1/mock cells. The cells for ChIP-seq were different culture from other sequences. For ChIP-qPCR and RT-qPCR of *a1* gene, yeast cells cultured in YPD medium were used.

### Construction of ChIP-seq library

ChIP was performed in the same way as ChIP-qPCR, except that culture condition was different and reverse crosslink was performed at 65℃ for overnight. Chromatin immunoprecipitation was performed using 2 µg anti-H3K9me2, 2 ug anti-H3K9me3, and 1 ug anti-HP1β (BMP002, MBL) antibodies. Libraries were constructed using KAPA Hyper Prep kit (KAPA BIOSYSTEMS) and xGen™ Stubby Adapter-UDI Primers (Integrated DNA Technologies). The concentration was quantified by KAPA Library quantification kit (KAPA BIOSYSTEMS). Sequencing was performed by HiSeq X (Illumina Inc.).

### RNA extraction

Cultured yeast cells in 5 ml YPD until O.D._600_=1 were collected and washed with distilled water. Cells were suspended in 200 µl AE buffer (50 mM sodium acetate, 10 mM EDTA, pH5.2) with 1% SDS. 200 µl phenol/chloroform (pH5.2) (nacalai tesque) were added to the suspension and incubated at 65℃ for 7 minutes. After centrifugation, aqueous layer was transferred to a new 1.5 ml tube, and RNA was purified by phenol/chloroform followed by ethanol precipitation.

### Construction of RNA-seq library

Purified RNA was treated with DNase I (New England Biolabs), and DNase treated RNA was purified by phenol/chloroform followed by ethanol precipitation. Libraries were constructed from 500 ng RNA using KAPA RNA HyperPrep kit (KAPA BIOSYSTEMS) and xGen™ Stubby Adapter-UDI Primers (Integrated DNA Technologies). Sequencing was performed by HiSeq X (Illumina Inc.).

### Construction of ATAC-seq library

1.5×10^7^ cells were incubated in 42.1 µl spheroplast buffer containing 1.26 µl 20 mg/ml zymolyase 20T for 30 minutes at room temperature. After centrifugation, cells were lysed in 100 µl ATAC buffer (10 mM Tris-HCl, 10 mM NaCl, 3 mM MgCl_2_, 0.1% NP40, pH7.4) and centrifuged. The pellet was suspended in 100 µl distilled water, and 18 µl was used for library construction. Construction of ATAC-seq samples was performed as previously described (Buenrostro, Giresi, Zaba et al., 2013). Nextera DNA library prep kit (illumina) and Nextera index kit (illumina) were used for the preparation. Sequencing was performed by HiSeq X (Illumina Inc.).

### Construction of Micro-C XL library

Construction of Micro-C XL library was performed as previously described (Hsieh et al., 2016). Details are in supprementary methods.

### ChIP-seq data analysis

Data analysis was performed using Galaxy platform (Afgan, Baker, Batut et al., 2018). Trimmed reads with FASTP (Chen, Zhou, Chen et al., 2018) were mapped to sacCer3 reference genome with Bowtie2 (Langmead, & Salzberg, 2012). PCR duplicates were removed with picard tools (http://broadinstitute.github.io/picard/). The data was visualized by Integrative Genomics Viewer. We also reanalyzed previously reported H3K9ac, H3K14ac, H3K18, H3K27ac and H3K36me3 ChIP-seq data (Weiner et al., 2015), which were deposited in the GSE61888.

### RNA-seq data analysis

Data analysis was performed using Galaxy platform. After trimming with FASTP, the quantification of transcripts was performed with Sailfish (Patro, Mount, & Kingsford, 2014). The differential analysis of the count data was performed by DESeq2 (Love, Huber, & Anders, 2014).

### ATAC-seq data analysis

Data analysis was performed using Galaxy platform. Trimmed reads with FASTP were mapped to sacCer3 reference genome with Bowtie2. After removing mitochondrial reads, PCR duplicates were removed with picard tools. Peak calling was performed with Genrich (https://github.com/jsh58/Genrich), and the data was visualized by Integrative Genomics Viewer.

### Correlation analysis between chromatin conformation and HP1β

The number of ATAC-seq reads in every 100 bp was counted by featureCounts (Liao, Smyth, & Shi, 2014), and each count was normalized by total read numbers. The normalized counts were compared between Suv39h1/mock and Suv39h1/HP1β (log_2_(normFC_HP1β+1_/normFC_mock+1_)). The amount of HP1β in the same 100bp region was counted by featureCounts using ChIP-seq data and normalized by input ChIP-seq data (normFC_HP1β_/ normFC_input_). The relationship between the amount of HP1β and chromatin conformational change was visualized by plotting these values.

### Micro-C XL data analysis

After mapping to sacCer3 by Bowtie2, the analysis was performed using HiCExplorer (Ramirez, Bhardwaj, Arrigoni et al., 2018; Wolff, Bhardwaj, Nothjunge et al., 2018; Wolff, Rabbani, Gilsbach et al., 2020). Micro-C XL matrices were built with hicBuildMatrix. The matrices were normalized to the smallest read count and corrected. TAD regions were extracted by hicFindTADs.

### RT-qPCR

1 µg purified RNA was treated with DNaseI (New England Biolabs) at 37℃ for 15 minutes, and DNaseI was deactivated by incubation at 70℃ for 10 minutes. Reverse transcription was carried out with Omniscript RT kit (QIAGEN). qPCR was performed using StepOnePlus (Thermo Fisher Scientific) with a1 primers. *ACT1* was used for internal control (Table S1)

### Mating assay

Yeast strains grown on YPD plate were plated on synthetic defined medium with *his4-*yeast strain and cultured for 2-3 days at 30℃ (Oki et al., 2004).

### Statistical analysis

Results were reported as mean ± SD. Statistical significance was determined by Student’s *t* test, Dunnett’s test or Tukey Kramer’s test. We evaluated that the value of p<0.05 was statistically significant.

## Data Availability

All sequencing data have been submitted to Gene Expression Omnibus under accession number GSE224641.

## Acknowledgements

We thank the staff of the Support Unit for Bio-Material Analysis (BMA) at the RIKEN Center for Brain Science (CBS) Research Resources Division (RRD) for NGS library construction, DNA sequencing. We would also like to thank Yota Murakami and Shinya Takahata for their valuable comments on this study, our colleagues at Shinkai laboratory for their support and valuable comments. This work was supported by the Japan Ministry of Education, Culture, Sports, Science and Technology Grant-in-Aid for Scientific Research (A) 18H03991, 22H00413 and Scientific Research on Innovative Areas 18H05530 (to Y.S.), Scientific Research (B) 17H03713, 20H03189 and Scientific Research on Innovative Areas 16H01315, 18H05532 (to J.N.), and the Special Postdoctoral Researcher (SPDR) Program of RIKEN (to K.F.).

