## Supplementary methods for "Chromatin landscape of budding yeast acquiring H3K9 methylation and its reader molecule HP1"

**Chromatin extraction**

 chromatin samples were used for western blotting of histone modifications. Cultured yeast cells on YPD plate were washed with cell wash buffer (50 mM Tris-HCl, 3 mM DTT, pH 7.5) and treated with zymolyase 20T (nacalai tesque) in spheroplast buffer (1M sorbitol, 25 mM Hepes-KOH, 5 mM MgCl_2_, pH 7.5, 1mM DTT) for 30 minutes. Cells were centrifuged and lysed in TritonX buffer (10 mM Tris-HCl, 50 mM KCl, 15 mM NaCl, 5 mM MgCl_2_, 1 mM CaCl_2_, 0.8% TritonX-100, 1 mM PMSF, pH7.5). After centrifugation, the supernatant was discarded, and the pellet was washed with high salt buffer for nuclear extraction (10 mM Hepes-KOH, 400 mM NaCl, 1 mM PMSF, pH7.5). After centrifugation, the pellet was used as an extracted chromatin.

**Western blot analysis**

Harvested yeast cells were suspended with extraction buffer (2 M NaOH, 7.5% β-mercaptoethanol) and rotated at room temperature for 5 minutes. After centrifugation, cells were washed by distilled water. The cells were lysed in SDS sample buffer (62.5 mM Tris-HCl, 5% sucrose, 2% SDS, 5% β-mercaptoethanol, pH 6.8) and incubated at 95 ℃ for 5 minutes. The lysate was centrifuged at 21,500 x g for 5min, and supernatant was separated by sodium dodecyl sulfate-polyacrylamide gel electrophoresis and transferred to a PVDF membrane. Blocking was performed in 2% bovine serum albumin for histone or in 5% skim milk for others for 1 hour. Primary antibodies were listed in Table S2. For histone proteins, IRDye^®^ 680RD Goat anti-Rabbit IgG Secondary Antibody (926-68071, LI-COR) and IRDye^®^ 800CW Goat anti-Mouse IgG Secondary Antibody (926-32210, LI-COR) were used as secondary antibodies, and they were detected and quantitated with Odyssey CLx (LI-COR Inc.). For other proteins, secondary antibodies conjugated with HRP were used. Detection was performed by chemiluminescence, and bands were visualized using an X-ray film.

**Immunostaining**

 Yeast cells were fixed in 4% paraformaldehyde at 37℃ for 15 minutes and washed with cell wash buffer.  Cell wall was digested by zymolyase 20T for 20 minutes and washed with PIPES/Sorb buffer (20 mM Pipes-KOH, 1.2 M Sorbitol, 1 mM MgCl_2_, pH6.8) for two times. The cells were plated on Poly-L-Lysine coated cover glass and stood for 15 minutes. The adhered cells on cover glass were permeabilized with 0.1% ToritonX-100 for 15 minutes at room temperature and blocked with 2% bovine serum albumin at 37℃ for 15 minutes. After blocking, cells were incubated with anti-G196 antibody (1:200) at 4℃ for overnight. Goat anti-Mouse IgG Alexa Fluor™ 488 (A11001, Thermo Fisher Scientific) was used as secondary antibody. 1 mg/ml DAPI was treated for 5 minutes at room temperature for nuclear staining. The immunofluorescence was imaged using FV3000 (Olympus).

**ChIP-qPCR**

Yeast cells were cultured in 50 ml YPD culture medium until OD_600_ = 0.8~1, and fixed in crosslink buffer (1% paraformaldehyde, 10 mM Hepes-KOH, 3 mM MgCl_2,_ pH7.5) for 20 minutes at room temperature. The fixed cells were quenched by 125 mM Glycine and washed with cell wash buffer. After centrifugation, cells walls were digested by 25 µg zymolyase 20T in spheroplast buffer until OD_600_=1/10. Cells were washed with PIPES/Sorb buffer and transferred into 1.5 ml tube by PBS. After centrifugation, cells were permeabilized with Hepes/TritonX-100 buffer (0.25% Triton X-100, 10 mM EDTA, 0.5 mM EGTA, 10 mM Hepes-KOH, 2 mM PMSF, pH 7.5) and centrifuged at 1,500 x g for 5 minutes. Pellet was washed with Hepes/NaCl buffer (200 mM NaCl, 1 mM EDTA, 0.5 mM EGTA, 10 mM Hepes-KOH, pH7.5) and lysed in 100 µl SDS lysis buffer (1% SDS, 10 mM EDTA, 50 mM Tris-HCl, pH8.0). The lysate was diluted by 375 µl IP dilution buffer (1.1% TritonX-100, 1.2 mM EDTA, 16.7 mM Tris-HCl, 167 mM NaCl, pH8.0) with PMSF, and sonicated by Bioruptor Pico (Diagenode) for 5 cycles of 30 seconds on/off. After centrifugation, supernatant was transferred into 1.5 ml tube with 690 ml IP dilution buffer and stored at -80℃.

 The lysate was incubated with protein A/G plus agarose (sc-2003, Santa Cruz) and a following primary antibody at 4℃ for overnight; 2 µg anti-H3K9me2 antibody (6D11), 4 µg anti-H3K9me3 antibody (2F3), 1 µg anti-HA antibody (014-21881, Fujifilm). The beads were washed in order of low salt buffer (0.1% SDS, 1% TritonX-100, 2 mM EDTA, 20 mM Tris-HCl pH8.0, 150 mM NaCl), high salt buffer (0.1% SDS, 1% TritonX-100, 2 mM EDTA, 20 mM Tris-HCl pH8.0, 500 mM NaCl), LiDW (0.25 M lithium chloride, 1% sodium deoxycholate, 10 mM Tris-HCl pH 8.0, 1% NP40, 1mM EDTA), and TE buffer (10 mM Tris-HCl, 1 mM EDTA, pH8.0). Reverse crosslink was performed in 200 µl SDS lysis buffer containing 200 mM NaCl at 95℃ for 15 minutes, and DNA was purified by phenol/chloroform followed by ethanol precipitation. Purified DNA was dissolved TE buffer containing 20 µg/ml RNase and incubated at 42℃ for 1 hour.

 qPCR was performed using StepOnePlus (Thermo Fisher Scientific). Primer list is Table S1.

**Construction of Micro-C XL library**

The procedures until zymolyase treatment were same as ChIP-seq protocol. Cells treated with zymolyase were washed with PBS and suspended in PBS with 3 mM disuccinimidyl glutarate (Thermo Fisher Scientific) for 45 minutes at room temperature and quenched by 667 mM Glycine. Chromatin was digested by micrococcal nuclease. The reaction was stopped by adding 4.67 mM EGTA/EDTA and incubating at 65℃ for 10 minutes.

Cells were suspended in NEBuffer 2.1 (New England Biolabs) with 5 U rSAP (New England Biolabs) and incubated at 37℃ for 45 minutes. rSAP was deactivated by incubation at 65℃ for 10 minutes. After inactivation, 2 mM ATP, 7.5 U Klenow Fragment (New England Biolabs), 15 U T4 polynucleotide kinase (New England Biolabs) and 3 mM DTT were added. After incubation at 37℃ for 30 minutes, dNTPs solution (70 µM biotin-dATP/biotin-dCTP (Jena Bioscience), 70 µM dGTP/dTTP (TOYOBO) in T4 DNA ligase buffer (New England Biolabs) with 1 mg/ml BSA) were added, and the suspension was incubated for 45 minutes at room temperature. After inactivation of enzymes by adding 3 mM EDTA and incubating at 65℃ for 20 minutes, cells were washed with NEBuffer 2 (New England Biolabs), and DNA was ligated by T4 DNA ligase (New England Biolabs) for 2.5 hours at room temperature. After exonuclease III treatment, cells were incubated at 55℃ for overnight with 200 µg/ml Proteinase K.

DNA was purified by phenol/chloroform followed by ethanol precipitation and incubated in 20 µg/ml RNase containing TE buffer at 30℃ for 30 minutes. Di-nucleosomal DNA was extracted by 3% Nusieve agarose DNA gel. Biotinylated DNA was purified by Dynabeads M-280 Streptavidin (Thermo Fisher Scientific).

 Libraries were constructed using KAPA Hyper Prep kit (KAPA BIOSYSTEMS) and xGen™ Stubby Adapter-UDI Primers (Integrated DNA Technologies). Sequencing was performed by HiSeq X (Illumina Inc.).
