## Supplementary table and figure for "Chromatin landscape of budding yeast acquiring H3K9 methylation and its reader molecule HP1"

### Table S1 Primers used in this study

| primer | Forward | Reverse |
|---|---|---|
| SAP1 | GTGAACCAGTCAGGGGGATG | AAAGGTGGAGTGCGACTCTG |
| THS1 | AGGTGATGCGACAAACGACT | TAGAGGCTTCCGCCAAGAAC |
| ACC1 | ACACATGGTTGTTGCCCTGA | GGCAAGAGTTGGATCAGGCT |
| preSAP1 | ACCCAAGCATCTTAGCCGAG | TTGTAGTTCTTGGGGAGCCG |
| preTHS1 | GACCTTTCCCACTCATCGCA | ACTTCCATAGGCGCTTGTCC |
| preACC1 | CGCTCATTGCCACTCTCTCT | TGTGCCAGAATACGACTGGG |
| a1 | GCCCAAAGGGAAAATCATCA | TCCTTGGAATTAAGGCTTTGCT |
| -3k | CGTTGGAAATCACTGGGAAA | GATGCATTGATGTGCCAAAG |
| -0.2k | TTGCAACAACTTCTTCTCCTCATC | ACCAGAGAGTGTAACAACAGAAGA |
| +2.4k | CGAACGATCCCCGTCCAAGTTATG | GCATTACGAAGATTCTCGATTCCGA |
| +3k | GCGATCAATGACGGTGCTCAATC | AAGAAAGCAACTCCCGTCACTT |
| ACT1 | TCGTTCCAATTTACGCTGGTT | CGGCCAAATCGATTCTCAA |

**Table S2 Antibodies list**

| Commercial Antibody | Cat. | Company | dilution |
|---|---|---|---|
| anti-Flag | F-1804 | Sigma Aldrich | 1:2000 |
| anti-G196 | R-G-001 | mAbProtein | 1:2000 |
| anti-HA | 11867423001 | Roche | 1:2000 |
| anti-SETDB1 | CP.10377 | Cell Applications | 1/2000 |
| anti-Suv39h1 | 8729 | Cell Signaling Technology | 1:2000 |
| anti-G9a | PP-A8620A-00 | R&D systems | 1:1000 |
| anti-GLP | PP-B0422-00 | R&D systems | 1:2000 |
| anti- histone H3 | 07-690 | EMD Millipore | 1:5000 |

| antibody | Clone ID | dilution |
|---|---|---|
| anti-H3K9me2 | 6D11 | 1:2000 |
| anti-H3K9me3 | 2F3 | 1:1000 |
| anti-H3K4me3 | 16H10 | 1:2000 |
| anti-H3K9ac | CMA305 | 1:2000 |
| anti-H3K27ac | 9E2H10 | 1:2000 |

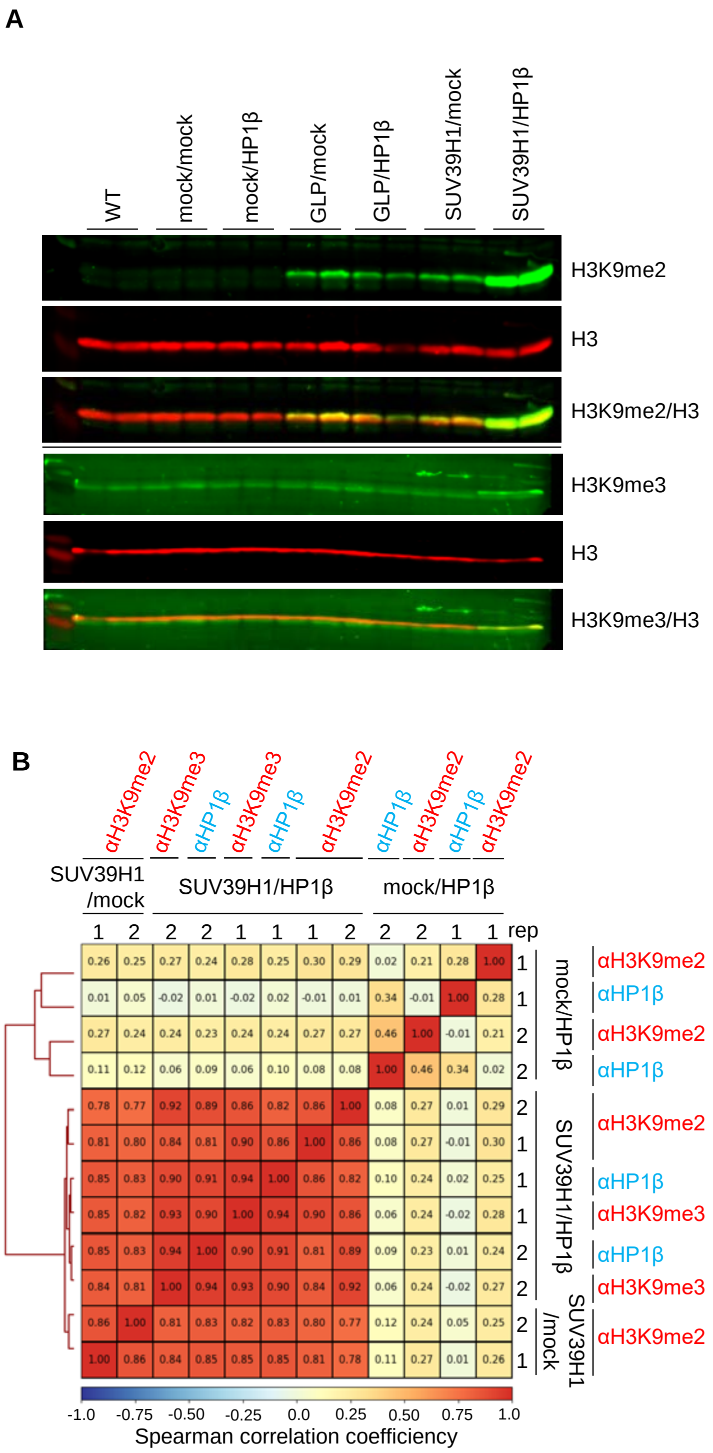


**Supplementary Figure 1**

**The correlation of H3K9 methylation and HP1β localizations in SUV39H1 strain.** (A) Cells dispensed from the cultured cells for ChIP-seq were lysed in SDS sample buffer and western blotting was performed using anti-H3K9me2, anti-H3K9me3, and anti-H3 antibodies. mock indicates G196 empty vector for H3K9 methyltransferases or HA empty vector for HP1. (B) The correlation between H3K9 methylation and HP1β in every strain (mock, SUV39H1) was determined by Spearman correlation heatmap. Both replication 1 and 2 data were shown.

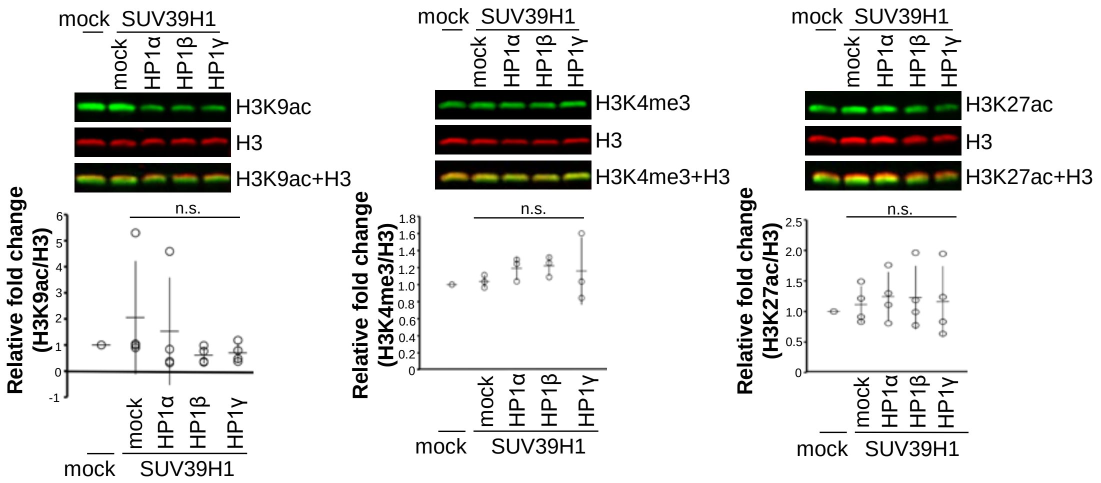


**Supplementary Figure 2**

**Histone modification in SUV39H1 strain.** The quantity of H3K9ac, H3K4me3, and H3K27ac in the same nuclei lysate as Figure 1E were determined by western blotting. The signal intensity was normalized by H3. Error bars indicate S.D. from three or four independent experiments. Statistical analysis was performed by Tukey Kramer’s test.

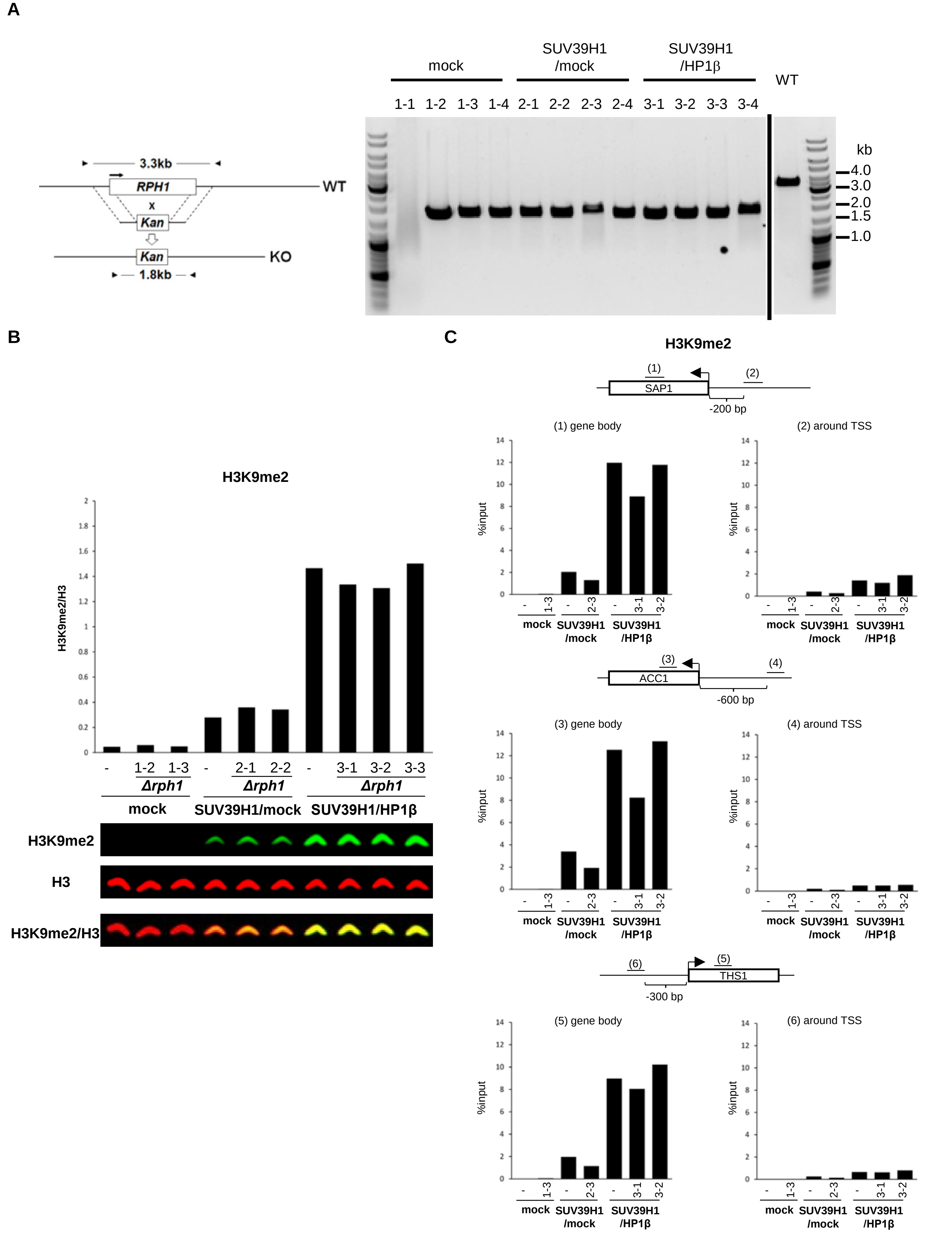


**Supplementary Figure 3**

**The effect of *rph1* deletion on H3K9 methylation level.** (A) *RPH1* gene was deleted in mock and SUV39H1 expressing strains using KanMX cassette. Deletion was confirmed by PCR (WT:3.3kb, KO:1.8kb), and some clones were used for the analysis. (B, C) The effect of *rph1* deletion in each strain (mock, SUV39H1/mock, SUV39H1/HP1β) on H3K9 di-methylation was determined by western blotting (B) and ChIP-qPCR (SAP1, ACC1, and THS1 gene loci) (C).

**A**

mock/mock vs SUV39H1/mock
(SUV39H1/mock)

| name | log2FoldChange | FDR |
|---|---|---|
| URA3 | 2.75648 | 4.75E-75 |
| STR3 | 1.1475 | 4.23E-08 |
| LEU2 | 1.0846 | 9.04E-07 |
| FIT3 | 0.78701 | 0.005922 |
| PHO89 | 0.71998 | 0.003932 |
| ENA2 | -1.11658 | 4.23E-08 |

SUV39H1/mock vs SUV39H1/HP1β
(HP1β/mock)

| name | log2FoldChange | FDR |
|---|---|---|
| ASG1 | 0.7905 | 0.005937 |
| YOR338W | 0.72304 | 0.007673 |
| BUD21 | 0.67843 | 0.028135 |
| RGS2 | 0.67583 | 0.024235 |
| TAH11 | 0.62142 | 0.028135 |
| LEE1 | -0.56028 | 0.01307 |
| GDE1 | -0.56247 | 0.030242 |
| HIS4 | -0.57421 | 0.001962 |
| ICY1 | -0.67707 | 0.047598 |
| MUP3 | -0.70193 | 0.028135 |
| ADH5 | -0.80003 | 0.002578 |
| FIT3 | -0.83278 | 0.001962 |
| MTD1 | -0.90422 | 0.0002 |
| STR3 | -1.11152 | 3.32E-07 |
| PHO89 | -1.36908 | 5.34E-15 |

**B**

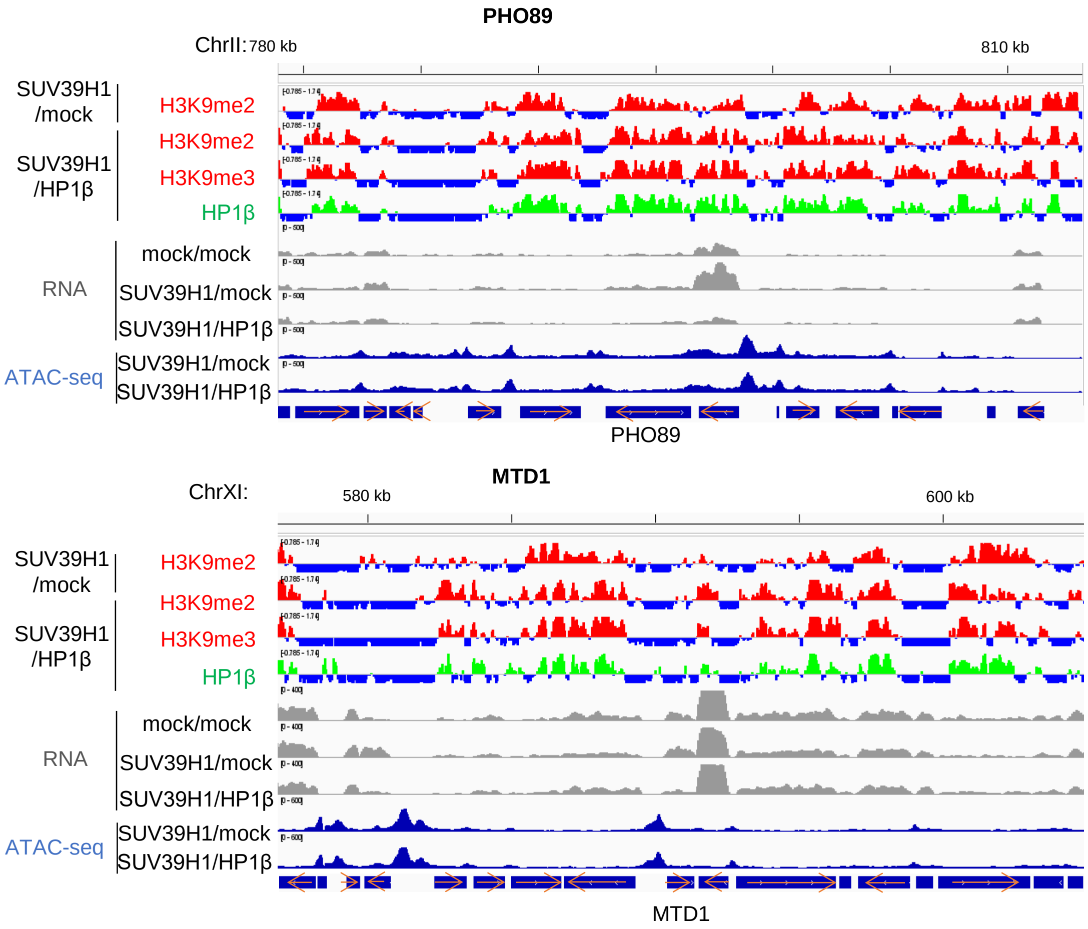


**Supplementary Figure 4**

**The effect of SUV39H1 and HP1β expression on mRNA expression level of budding yeast.** (A) The name of genes with significantly different expression level (FDR<0.05) were shown in table. The left table indicates the effect of SUV39H1 expression (SUV39H1/mock vs mock/mock). The right table indicates the effect of HP1β expression (SUV39H1/HP1β vs SUV39H1/mock). (B) RNA expression level in *PHO89* and *MTD1* gene loci were viewed in Integrative Genomics Viewer with ChIP-seq data (H3K9me2/H3K9me3/HP1β) and ATAC-seq data.

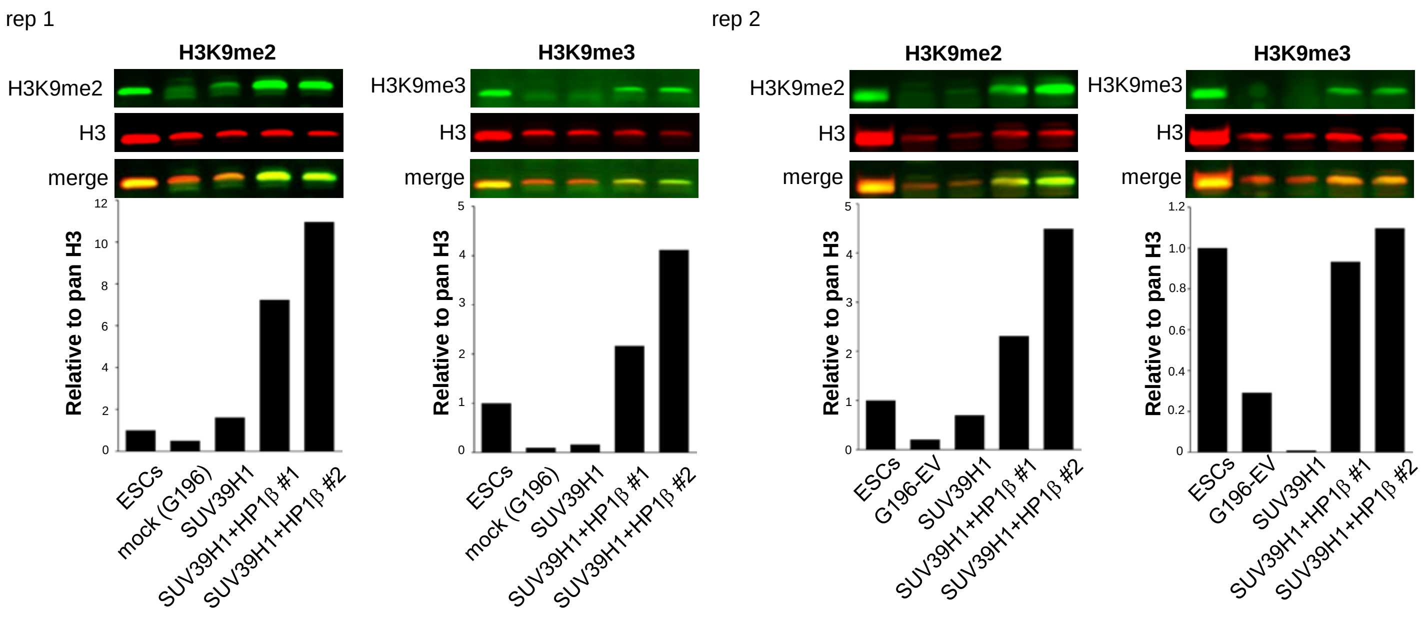


**Supplementary Figure 5**

**Comparison of H3K9me2 and 3 levels between mouse ESCs and budding yeast expressing SUV39H1 or SUV39H1+HP1β.** H3K9 di-, tri- methylation level in SUV39H1 strains was compared to that in mouse ES cells by western blotting. The signal intensity was normalized by H3.

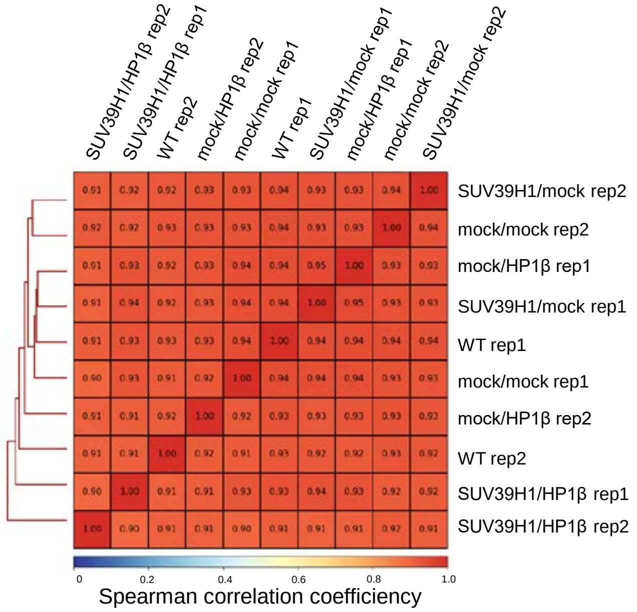


**Supplementary Figure 6**

**Correlation efficiency of ATAC-seq peaks.** The spearman correlation heatmap of every ATAC-seq mapping data was calculated. Both replication 1 and 2 were shown.

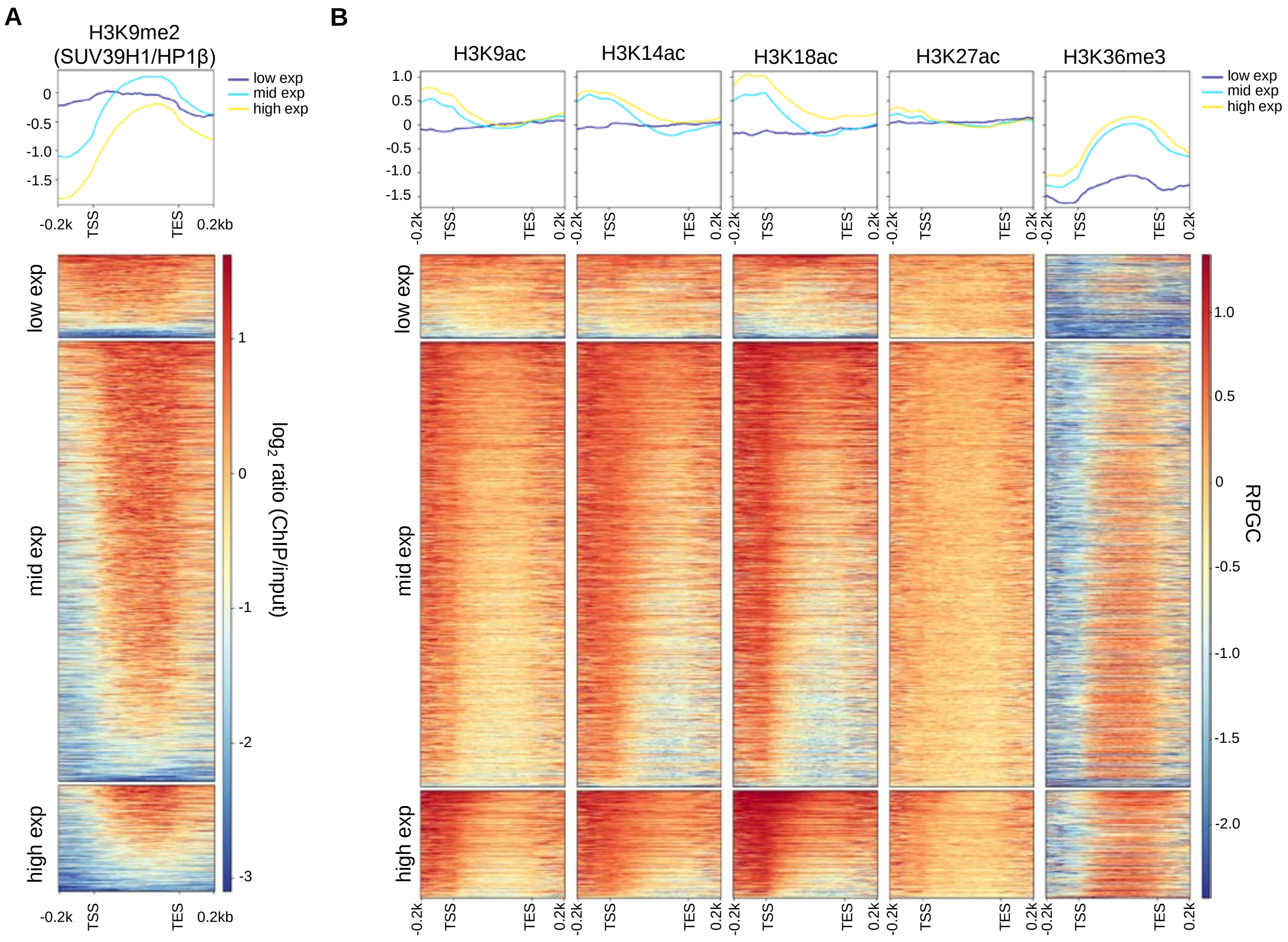


**Supplementary Figure 7**

**The region of histone modifications by gene expression level.** The genes were divided into three groups by the TPM values of WT rep1 RNA-seq data (low exp: <5 TPM, mid exp: 5-80 TPM, high exp: >80 TPM). The region of H3K9me2 (A) and the region of some histone modifications (H3K9ac/H3K14ac/H3K18ac/H3K27ac/H3K36me3) (B) in each group were ploted. ChIP-seq data from SUV39H1/HP1β rep1 was used for (A), and public ChIP-seq data was used for (B).

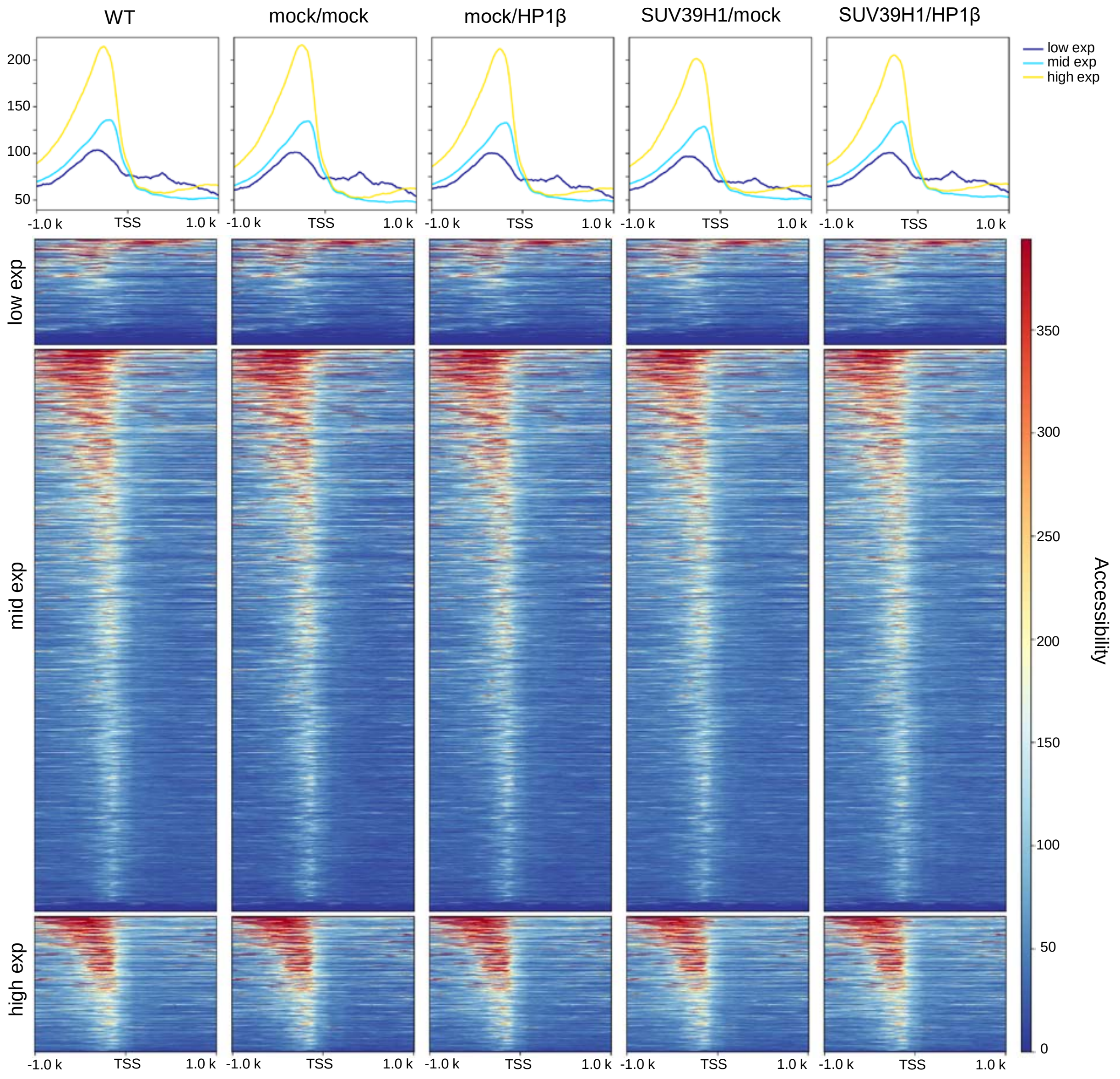


**Supplementary Figure 8**

**Genome accessibility by gene expression level.** The genes were divided into three groups by the TPM values of WT rep1 RNA-seq data (low exp: <5 TPM, mid exp: 5-80 TPM, high exp: >80 TPM). The genome accessibilities by gene expression level in each strain (WT, mock/mock, mock/ HP1β, SUV39H1/mock, SUV39H1/HP1β) were shown as heatmap.

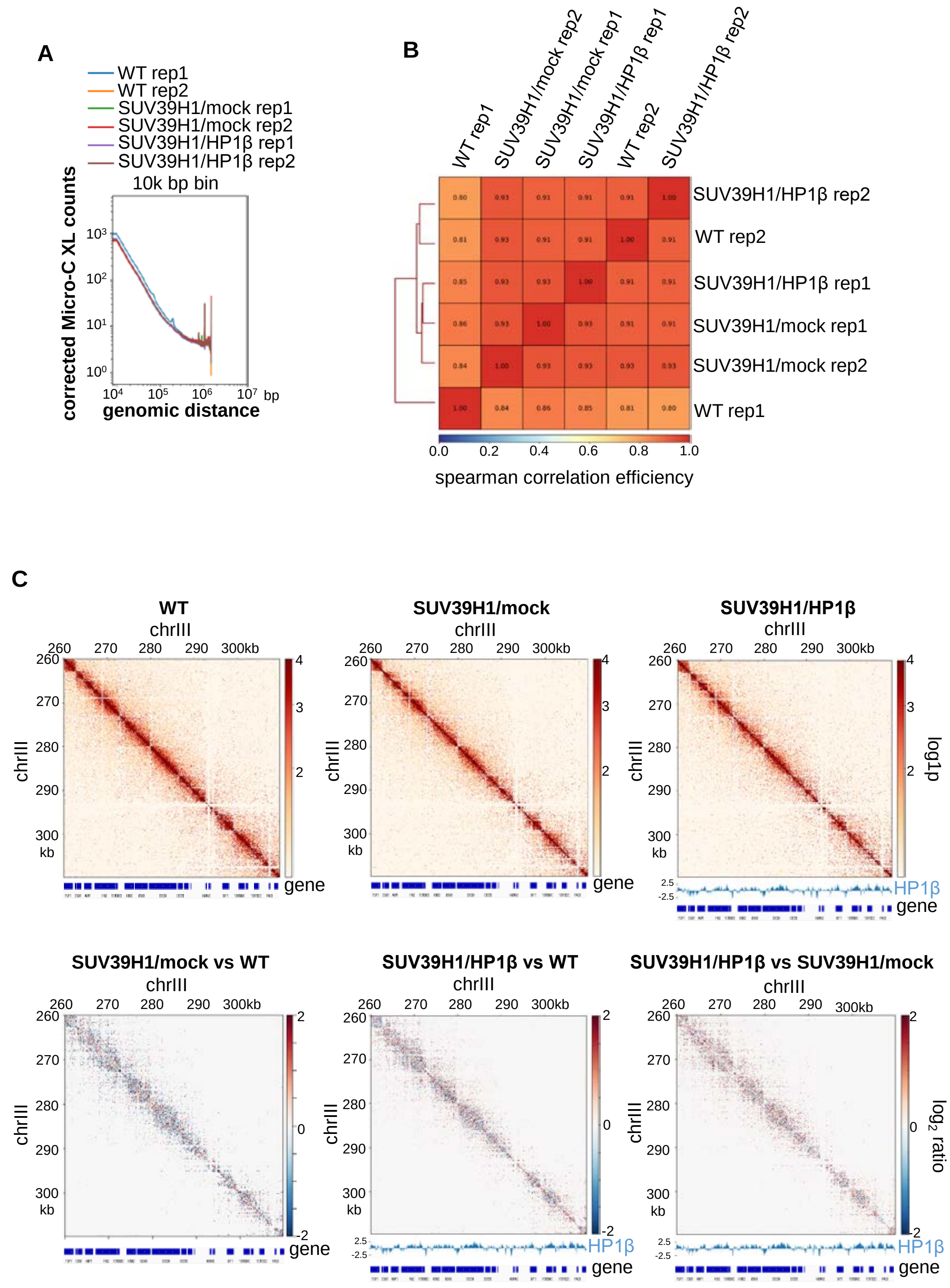


**Supplementary Figure 9**

**Comparison of TAD like structure in WT, SUV39H1/mock, and SUV39H1/HP1β strains.** (A) Genome contact frequency in WT, SUV39H1/mock, and SUV39H1/HP1β strains were compared using 10k bp windows. Both replication 1 and 2 were shown. (B) TAD regions in 100 bp bin contact matrix was identified, and the spearman correlation heatmap was plotted. Both replication 1 and 2 were shown. (C) Genome contact heatmap in chromosome III 260kb-310kb at 100 bp resolution. Top panels show each contact heatmaps, and bottom panels show compared contact heatmaps. Merged matrix data were used, and SUV39H1/HP1β data was shown with ChIP-seq data of HP1β from SUV39H1/HP1β strain replication 1.

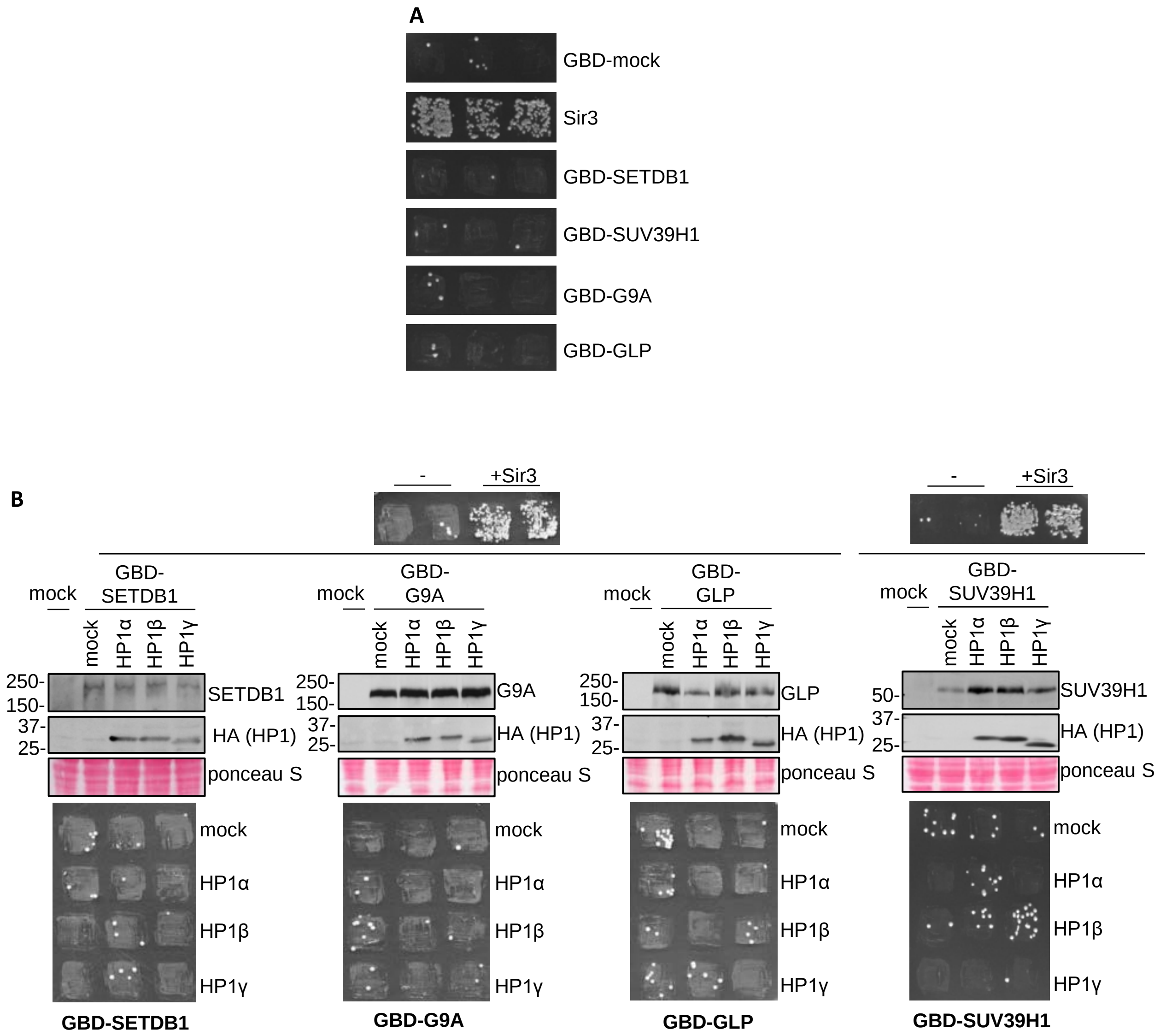


**Supplementary Figure 10**

**Mating assay for a1 gene transcriptional analysis.** (A) Each GBD-H3K9 methyltransferase or Sir3 expressing ROY2042*Δsir3* strain grown on YPD plate was mixed with *his4*- strain and cultured on synthetic defined medium plate. Mock indicates GBD empty vector. (B) HA-empty vector, HA-HP1α, HA-HP1β, or HA-HP1γ was co-expressed in each GBD-H3K9 methyltransferase expressing ROY2042*Δsir3* strain. The expression of H3K9 methyltransferases and HP1 was determined by western blotting (top). mock indicates GBD empty vector expressing strain. These strains were mixed with *his4*- strain and cultured on synthetic defined medium plate (bottom).
